# Megabase-scale detachment of genome-nuclear lamina interactions is required for efficient repair of double stranded breaks

**DOI:** 10.1101/2025.07.03.662902

**Authors:** Xabier Vergara, Anna G. Manjón, Marco Novais-Cruz, Marcel de Haas, Ayoub Ouchene, Tom van Schaik, Stefano G. Manzo, Richard Cardoso da Silva, Tuncay Baubec, Bas van Steensel, René H. Medema

## Abstract

Lamina-associated domains (LADs) are large genomic regions that contact the nuclear lamina (NL). Double-strand breaks (DSBs) in LADs are known to be repaired more slowly and with different pathway preferences compared to other chromatin contexts. However, little is known about the chromatin changes at LADs that occur during DSB repair. Here, we report that a single DSB inside a LAD can cause detachment from the NL over several megabases. This profound spatial rearrangement is transient and reverts within 48 hours. Preventing this detachment slows down repair kinetics and renders repair incomplete, indicating that NL detachment is required for efficient repair of DSBs in LADs. NL detachment is dependent on γH2AX and ATM, while it is antagonized by DNAPK_cs_ activity. Remarkably, γH2AX also antagonizes NL interactions at chromosome ends. Taken together, our data indicate that γH2AX accumulation in LADs induces large scale rewiring of genome-NL interactions, allowing for efficient repair of DSBs.

## INTRODUCTION

DNA double-strand breaks (DSBs) occur daily in human cells, and failure to repair them leads to genomic instability^1^. In human cells, multiple pathways repair DSBs, including non-homologous end-joining (NHEJ), homologous recombination (HR), microhomology-mediated end-joining (MMEJ) and single strand annealing (SSA) (reviewed in ^2,3^). NHEJ directly ligates the broken ends with very little DNA end-processing^4^. In contrast, HR, MMEJ and SSA require DNA end-resection and recombination between homologous sequences to ligate the broken ends^2^. DSBs also trigger chromatin changes that promote binding of DNA repair proteins and DNA repair efficiency (reviewed in ^5^).

One of the best characterized DSB-induced chromatin modifications is the phosphorylation of H2AX on serine 139 (γH2AX)^6^. Several protein kinases (e.g. ATM, DNAPK_cs_ or ATR) phosphorylate H2AX over megabases around the break site^7–9^. The resulting γH2AX domain acts as a binding platform for the adaptor protein MDC1^10,11^. Although recruitment of several DNA repair proteins to the break site depends on γH2AX-mediated binding of MDC1, cells lacking H2AX do not exhibit severe DNA repair deficiencies^12^. Some studies have shown that γH2AX fine-tunes the efficiency of DNA repair pathways, which is critical for class switch recombination^13,14^ and the repair of a subset of DSBs dependent on ATM and the Artemis nuclease^15^. However, the exact requirements for H2AX phosphorylation in the repair of DSBs remain poorly understood.

Besides acting as binding platforms for DNA repair proteins, DSB-induced chromatin changes have been shown to play a critical role in the repair of DSBs that are otherwise difficult to repair. These breaks tend to localize within heterochromatic areas of the genome^16^, which are marked by the histone modification H3K9me2/3 and are transcriptionally inactive(reviewed in ^17^). The repair of these breaks requires the removal of heterochromatin-associated proteins, such as H1 linker histone^18,19^, KAP1^20^, and/or HP1^21^. Collectively, DSB-induced chromatin changes in H3K9me2/3 heterochromatin promote the removal of heterochromatin associated proteins, and the efficient repair of these DSBs. In *Drosophila* cells, DSB repair in this type of heterochromatin triggers the removal of H3K9me2/3^22^. However, H3K9me2/3 levels were reported to remain unchanged upon a DSB in human cells^23^.

In addition to triggering the displacement of heterochromatin-associated proteins, DSBs in heterochromatin also exhibit directed movement towards the outside of the heterochromatic domain^24–29^. DSBs in euchromatin also exhibit increased movement rates during repair, but this appears to be driven by clustering of DSBs and/or homology search^30,31^. Re-localization of heterochromatic DSBs towards the outer edge of the heterochromatic domain enables the binding of DNA repair proteins such Rad51 and BRCA2 that cannot access the interior of the heterochromatic domain^28^. The re-localization depends on the Smc5/6 complex^25^ and the DNA end-resection machinery^28^. In most organisms the movement is limited to the vicinity of the heterochromatic domain^24,25,28^, but DSBs in *Drosophila* chromocenters, which are heterochromatic domains found in the nuclear interior, exhibit long-range movement towards the nuclear periphery^27,32^. This long-range movement in *Drosophila* cells is dependent on nuclear pore complex proteins, nuclear actin filaments, and myosins^33^. Regardless of its range, movement of heterochromatic DSBs is thought to promote repair by enabling binding of HR proteins that cannot access the heterochromatic domain.

In most metazoan cells, heterochromatin of various compositions is located at the nuclear lamina (NL) in domains that are named lamina-associated domains (LADs). LADs collectively take up to 40% of the genome in most human cells^34^. The few studies that induced DSBs in LADs showed that these breaks are repaired more slowly and with a higher MMEJ frequency than breaks in euchromatin^35,36^. Moreover, ATM and SMC5/6, which promote DSB repair in H3K9 di-or trimethylated heterochromatin, regulate DNA repair pathway balance in LADs^37^. However, it remains unclear whether extensive chromatin changes and/or DSB movement outside the LAD occurs during DSB repair in LADs. A prior study in which a DSB was induced in a locus that was artificially tethered to the NL failed to detect movement away from the NL^36^. Whether DSBs in LADs move towards the nuclear interior in an untethered system is currently unknown.

Here, we report that genome-NL interactions in human cells are transiently lost during DSB repair. Mapping of genome-NL interactions^38^ upon Cas9-induced breaks in LADs showed that megabases of the genome around the DSB detach from the NL. This detachment could also be visualized by microscopy. Furthermore, preventing the detachment by forced tethering leads to slower repair kinetics and a larger fraction of unrepaired DSBs, indicating that detachment from the NL is required for efficient repair of DSBs in LADs. We find that NL detachment requires the presence of H2AX and is partially suppressed when ATM is inhibited, suggesting that the detachment is, at least in part, driven by H2AX phosphorylation. Moreover, we discovered that γH2AX accumulation in subtelomeric regions of the chromosome also influences the NL interaction of these regions. Taken together, this study shows that (i) megabases of genome-NL interactions are rewired during DSB repair in a γH2AX-dependent manner, (ii) detachment from the NL promotes efficient repair of DSBs in LADs and (iii) γH2AX antagonizes NL interactions around DSBs and at chromosome ends.

## RESULTS

### Efficient induction of a double strand break in LADs

To study how DSBs affect NL interactions, we used Cas9 to induce DSBs at defined positions in LADs in immortalized non-transformed human RPE-1 cells^41,42^. For this we initially designed a synthetic guide RNA (gRNA1) (**Table 1**), which targets the center of a 7.1 Mb long LAD (LAD1, **Figure S1A**). LAD1 is conserved across different cell types, including RPE-1 and K562 cells (**Figure S1B**) and is surrounded by three smaller LADs in RPE-1 cells (∼ 1 Mb each). In RPE-1 cells, most of LAD1 overlaps with H3K9me3 and late replicated chromatin but does not contain H3K27me3 domains (**Figure S1A**). Thus, LAD1 recapitulates typical features of LADs in RPE-1 cells^34^. To test whether gRNA1 induces DSBs at a high frequency, we first measured the editing efficiency in LAD1 using TIDE^43^ at 48 hours after transfection. This analysis showed that gRNA1 produces high frequencies of short indels (∼ 60%) in LAD1 in RPE-1 cells (**Figure S1C)**, indicating that Cas9 can efficiently introduce breaks in LADs in this setting. Next, we determined the fraction of broken DNA at different timepoints after gRNA1 transfection using ligation-mediated PCR (LM-PCR**, See methods, Figure S1D-E**)^44^. We found a maximum of ∼ 10% of gRNA1 target sites to be broken simultaneously, indicating that cutting occurs in an asynchronous manner. The fraction of broken DNA decreased after 10 hours, reaching background levels at 48 hours. These data suggest that the DSBs induced by gRNA1 in RPE-1 cells are generally repaired within 48 hours (**Figure S1F**). We concluded that this experimental setting is well-suited to study how a DSB in a LAD affects NL interactions. It is important to note that we never detect more than ∼10% of broken fraction at any given timepoint, which may lead to a substantial underestimation of the effects described below.

### DSBs trigger local rewiring of NL associations across several megabases

Next, we mapped NL interactions across the entire genome using pA-DamID of LMNB2, a method that is particularly suited to generate detailed maps of NL interactions with high temporal resolution^38^. We initially analyzed NL interactions at 10 hours after gRNA1 transfection. To test whether a single DSB perturbs NL interactions globally, we first analyzed the genome-wide correlation between gRNA1 and mock-transfected samples. We observed very high correlation (Pearson’s R = 0.97, **Figure S1G**), indicating that a single DSB does not alter NL interactions globally.

To study *local* NL interaction changes, we then focused on a 15 Mb window around the gRNA1 target site (chr7:80.2-95.2Mb, **Figure S1A**) and visualized NL interaction data in two ways. First, we plotted pA-DamID tracks from mock (control, orange) and gRNA1-transfected (purple) cells (**Figure 1A**, bottom panel). Second, we plotted a multi-scale domainogram^39,40^ to visualize regions that significantly gained (purple) or lost NL interactions (orange) at different window sizes (**Figure 1A**, top panel).

**Figure 1:**
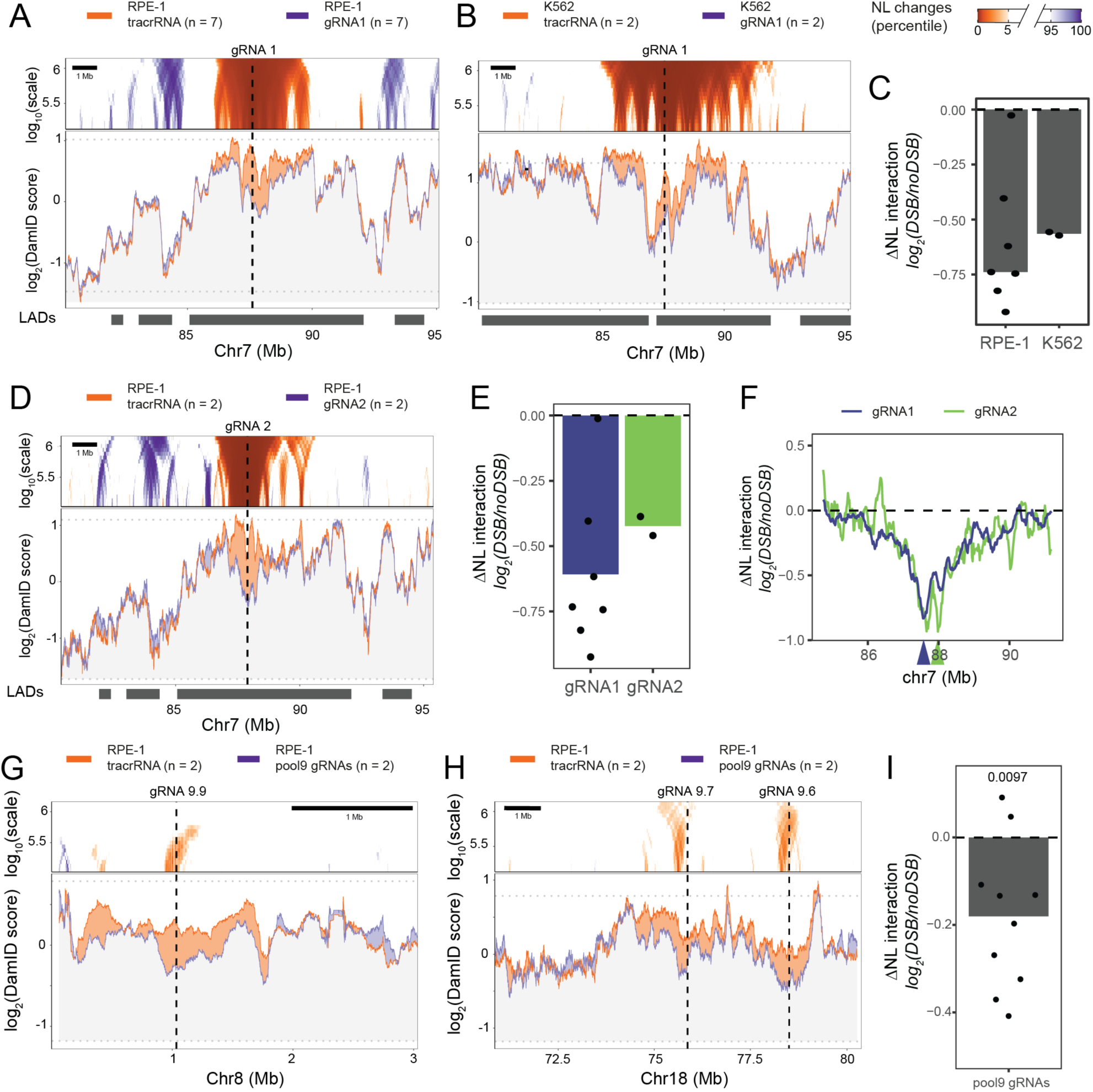
DSBs trigger rewiring of genome-NL interactions. (**A**) Comparison of NL interactions in Cas9-expressing RPE-1 cells transfected with gRNA1 (crRNA::tracrRNA duplex) to induce a single DSB (experimental, purple) or with tracrRNA alone (control, orange), in a window of 15 Mb (chr7:80.2-95.2 Mb) around the break site (vertical dashed line). Bottom panel: log_2_(LMNB1/Dam) pA-DamID tracks as two lines and the shading between them represents the color of the highest one. Horizontal dotted grey lines represent the 95% confidence interval of the genome-wide NL interaction scores, and the scale bar shows the size of the smoothing window. Top panel: “domainogram” visualization ^39,40^ showing significant NL changes at different bin sizes (vertical axis). For each bin size, regions with the largest 5% NL interaction gains (purple) or losses (orange) genome-wide are highlighted. (**B**) Same as Figure 1A for K562 cells. (**C**) Quantification of the differential NL interaction (ΔNL interaction) scores in a 1 Mb window around the break site calculated as the log_2_(gRNA1/tracrRNA only) for each replicate individually (dots) and all replicates combined (grey bar) in RPE-1 and K562 cells. (**D**) Domainogram visualization, as described in Figure 1A, of NL interaction changes 10 hours after gRNA2 (purple, n = 2) and tracrRNA only (orange, n = 2) transfection. ΔNL interaction scores, (**E**) as described in Figure 1C. (**F**) Direct comparison of gRNA1-versus gRNA2-induced changes as a differential track over LAD1 (chr7:84.74-91.2 Mb), 10 hours after gRNA2 transfection. (**G, H**) NL interaction changes 10 hours after pool 9 (n = 2) and tracrRNA only (n = 2) transfection, plotted 83 as described in Figure 1A, in a ∼ 3 Mb window around gRNA 9.9 (chr8:1,033,674) (**G**) and a ∼ 9 Mb window around gRNA 9.7 (chr18:75,860,915) and gRNA 9.6 (chr18:78,506,575) (**H**). (**I**) ΔNL interaction score, as calculated in Figure 1C, for each gRNA in pool 9 (n = 10). Each dot represents a single gRNA target site, and the grey bar is the 87 average across all 10 gRNA sites.

Strikingly, the single DSB in LAD1 caused a region of ∼3.4 Mb around the cut site to detach significantly from the NL in RPE-1 cells (**Figure 1A**). This detachment is much more extensive than the changes in NL contacts triggered by activation of a LAD-resident gene, which typically are limited to 50-100 kb of flanking sequence^40^. To exclude cell type-specific effects, we also transfected gRNA1 in K562 cells. We identified similar megabase-scale (6.3 Mb) NL detachment in these cells (**Figure 1B & S1G, Pearson’s R = 0.9**), although the exact pattern of losses differed. In K562 cells, NL interaction losses spread to neighboring LADs, indicating that DSB-induced NL rewiring is not confined by LAD borders, at least in this setting (**Figure 1B**). Quantitative analysis revealed that a single DSB can induce up to 40% reduction in NL interactions of the 1 Mb genomic region surrounding the DSB in both cell lines (**Figure 1C, p-value = 1.84**×10^-3^). This is most likely an underestimation of the actual magnitude of the effect, because of the asynchronous cutting by Cas9.

The domainogram also revealed more subtle but significant gains of NL interactions in RPE-1 cells. The regions that gain NL interaction coincide with the inter-LAD regions on each side of LAD1 in RPE-1 cells (**Figure 1A**). We speculate that these regions normally make few contacts with the NL, but occupy the surface of the NL that was vacated by the detached LAD (**Figure S1H**). Similar putative compensatory movements have been previously reported at a smaller scale upon changes in gene activity^40^. However, we cannot detect such gains in K562 cells (**Figure 1B**). Thus, gains of NL interaction might be influenced by the precise (cell type-dependent) LAD architecture of the locus.

### DSBs induce NL detachment across multiple LADs

Next, we studied whether the observed changes apply more generally to DSBs in LADs. To test this, we measured NL interactions after transfection of a different gRNA targeting LAD1, ∼300 kb away from gRNA1 (gRNA2, sequence in **Table 1**). Mapping of NL interactions after gRNA2 transfection revealed a near-identical NL detachment as observed for gRNA1 (**Figure 1D-E**). Quantitative analysis of the changes in NL interactions across LAD1 (chr7:84.74-91.2 Mb) showed that NL detachment induced by both gRNAs is highly similar (**Figure 1F**), with only subtle differences close to their target sites (arrowheads in **Figure 1F**).

To test whether NL detachment is conserved across other LADs, we mapped NL interactions after transfection of a pool of 10 gRNAs targeting non-repetitive heterochromatin domains (sequences in **Table 1**)^42^. These domains were often located in pericentromeric or sub-telomeric heterochromatic regions (**Figure S2A**) and normally contact the NL with modest frequencies (**Figure S2B**). Because we transfected ten gRNAs simultaneously, we first confirmed that genome-wide NL interactions remained unaltered (**Figure S2C**, Pearson’s R = 0.88). In contrast, three of the targeted loci showed megabase-scale loss of NL contacts (**Figure 1G-H & S2D-J**), and for the ten DSBs collectively this loss was significant (**Figure 1I**, p = 0.0097). We note that previous indel measurements of gRNAs 9.4, 9.5 and 9.10 indicated that the editing efficiencies of were substantially lower than for gRNA1 (on average 0.2 vs. 0.6), and gRNA 9.3 showed no detectable cutting^42^. Thus, this result is likely to underestimate the effects of DSBs on NL interactions, although we cannot rule out that some LADs are refractory to the detachment effect.

### NL detachment depends on break formation and is transient

We considered that the mere binding of Cas9, rather than its cutting activity, could directly lead to detachment of LAD1 from the NL. To rule this out, we measured NL interaction after transfection of a 15-nt (instead of the normal 20-nt) variant of gRNA1 in RPE-1 cells (**Table 1**). gRNAs of 15-nt can trigger the recruitment of Cas9 to the target site but fail to activate its nucleolytic activity^45^. In the absence of a DSB (**Figure S3A**), we did not observe NL interaction losses around the gRNA1 target site (**Figure 2A & S3B**). However, we detected subtle gains of NL interactions in the inter-LAD sequence next to LAD1 (**Figure S3B**). Possibly, this inter-LAD region is relatively unstable and particularly sensitive to subtle changes in the local chromatin state. Nevertheless, we conclude that the profound NL interaction losses are not caused by mere recruitment of Cas9 protein to LAD1 but require induction of a DSB.

**Figure 2:**
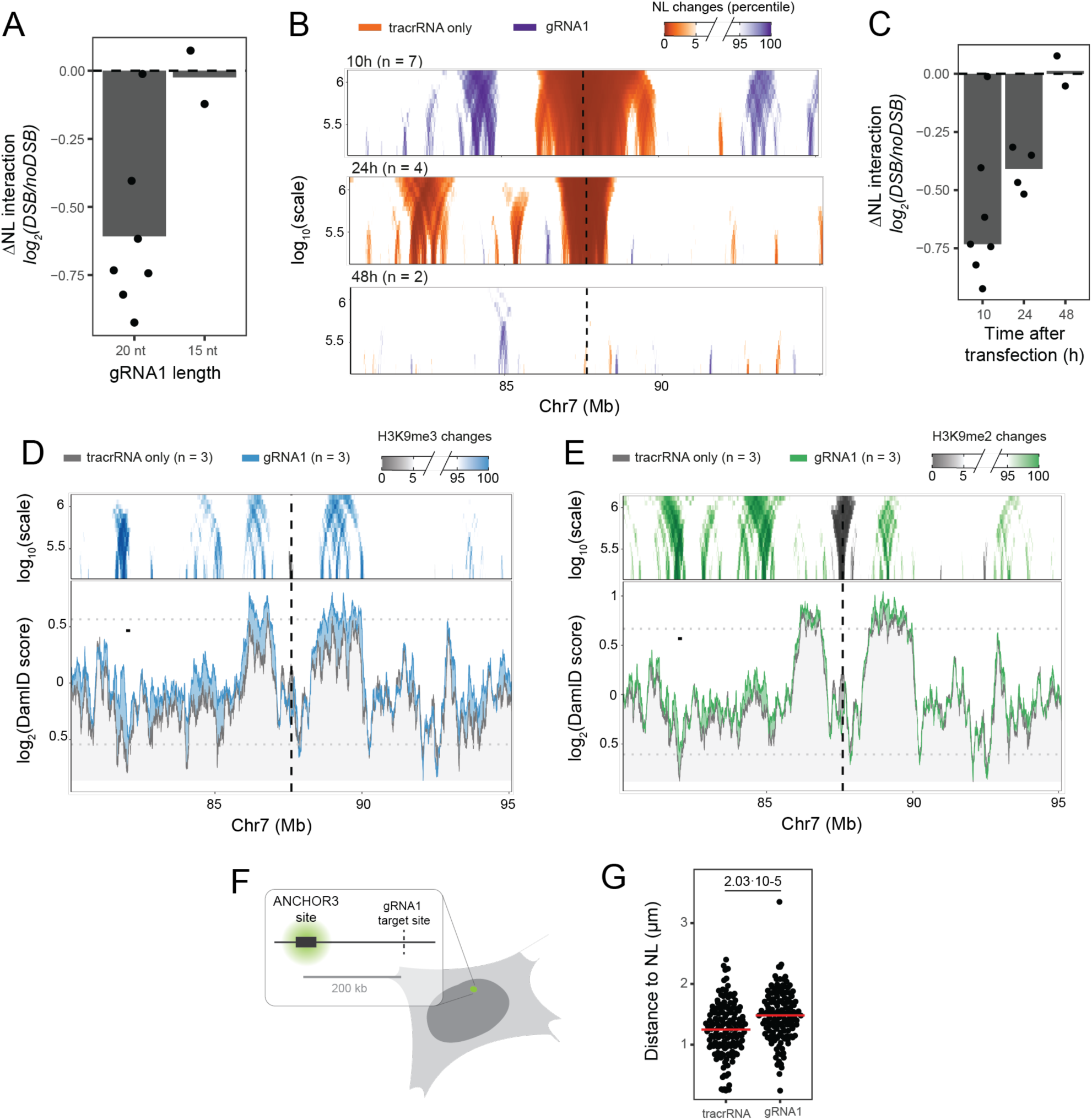
NL detachment requires Cas9-induced break formation, is transient and does not lead to H3K9 methylation changes. (**A**) ΔNL interaction scores, as calculated in Figure 1C, in RPE-1 cells transfected with a 20 nucleotide long (n = 7, same data as in Figure 1C) or 15 nucleotide long (n = 2) gRNA1. (**B**) Domainogram visualization, as described in Figure 1A, and (**C**) differential scores, as calculated in Figure 1C, of NL interactions 10 hours (n = 7, same data as in Figure 1A**&C**), 24 hours (n = 4) and 48 hours (n = 2) after gRNA1 transfection in RPE-1 cells. (**D**) H3K9me3 (in blue, n = 3) and (**E**) H3K9me2 (in green, n =3) score changes 10 hours after gRNA1 transfection in RPE-1 cells visualized as described in Figure 1A. (**F**) Cartoon describing how the RPE-1 ANCHOR3 cell line was built. (**G**) Shortest distance to the NL (µm) per RPE-1 ANCHOR3 cell with an OR3 focus 10 hours after tracrRNA only or gRNA1 transfection in four biological replicates. The red line represents the mean distance for each condition to the NL. p-values were calculated using a two-sided Student’s t-test.

Next, we studied the kinetics of NL detachment of LAD1 after DSB induction. For this we measured NL interactions at later timepoints (24 and 48 hours) (**Figure 2B & S3C-D**). Already after 24 hours, NL interactions were partially recovered, but it should be noted that at 24 hours we still detect a significant fraction of broken DNA. NL interactions of LAD1 were fully restored 48 hours after transfection (**Figure 2C**). These data indicate that NL detachment is not permanent, and that the NL interactions recover over time.

### NL detachment is not accompanied by loss of heterochromatic histone modifications

To test if NL detachment is accompanied by changes in other heterochromatic features, we mapped H3K9me2, H3K9me3 and H3K27me3 in RPE-1 cells (n = 3). We did not observe significant megabase-scale losses of any of these features. We only detected local losses of H3K9me2/3 around the target site (∼ 100 kb) and local gains scattered throughout LAD1 (**Figure 2D-E**). Both local removal and maintenance of H3K9me2/3 have been previously reported to occur upon DSB induction in different systems^22,23^. Our data provide a possible rationale for these discrepancies, since we find that H3K9me2/me3 is lost close to the break site but gained at other regions which are more remote from the break. We did not observe any significant change of H3K27me3 (**Figure S3E**), but we note that baseline H3K27me3 levels in LAD1 are low (**Figure S1A**). In summary, these results indicate that the large-scale NL interaction loss is not accompanied by similar changes in the most prevalent heterochromatic histone marks.

### NL detachment of DSBs in LADs can be visualized by microscopy

Because pA-DamID probes molecular contact frequency rather than physical distance, we wondered how far the DNA around the DSB moved away from the NL. To address this, we used an engineered RPE-1 cell line that contains an ANCH sequence in LAD1 (200 kb away from gRNA1) and expresses the ANCH-binding fluorescent protein OR3-GFP. This system allows the visualization of the tagged locus by confocal microscopy^46^ (**Figure 2F**). We then measured the distance of the OR3-GFP focus to the NL, with and without induction of the DSB 10 hours after transfection. In this experiment, we found that LAD1 was on average ∼250 nm further from the NL in gRNA1-transfected cells than in control-transfected cells (**Figure 2G & S3F**, p-value = 2.03×10^-5^). We note that this is likely to be an underestimate, because (i) we could not discern cells with a break in LAD1 from cells with an intact LAD1 at the moment of visualization; and (ii) accuracy of the distance measurements was compromised by the highly flattened morphology of the nuclei in combination with the limited z-resolution of confocal microscopy. Despite these limitations, these results confirm that DSB induction leads to LAD1 movement away from the NL, on average over several hundred nm.

### NL detachment correlates with γH2AX accumulation around DSBs

Previous studies have shown that γH2AX can spread across megabases flanking the break site^5,47^. To test whether the NL detachment correlates with γH2AX accumulation, we mapped presence of γH2AX across the genome at 10 hours after gRNA1 transfection. While mapping of γH2AX has been previously done by chromatin immunoprecipitation ^47^, we decided to use pA-DamID (which has been shown to reliably map other histone modifications ^38^) for the most direct comparison to our NL interaction data. The results confirmed that γH2AX forms a large domain around the break site in LAD1 (**Figure S4A**), with significant increases over a nearly 10 Mb long region around the break site (**Figure 3A-B**). Quantification of individual replicates showed that the γH2AX accumulation is reproducibly detected (**Figure 3C**).

**Figure 3:**
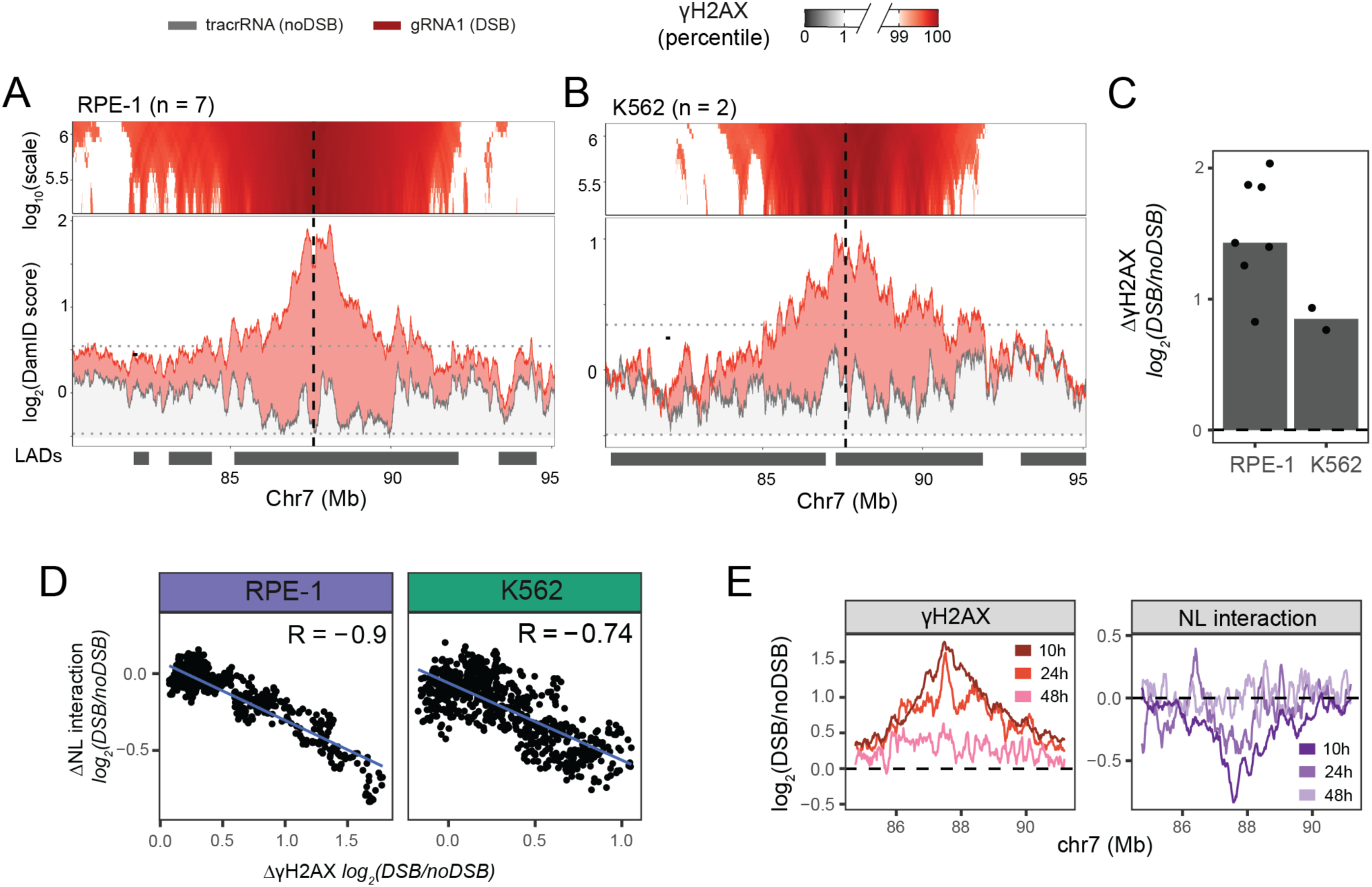
NL detachment correlates with gH2AX accumulation around DSBs. (**A**) gH2AX changes 10 hours after gRNA1 (red, n = 7) or tracrRNA only (grey, n = 7) transfection in RPE-1 cells. The top panel shows the “domainogram” visualization^39,40^ that highlights significant gH2AX changes at different bin sizes (vertical axis). For each bin size, regions with the largest 1% NL interaction gains (purple) or losses (orange) genome-wide are highlighted. (**B**) Same as in Figure 3A for K562 cells. (**C**) Quantification of the differential H2AX phosphorylation (ΔgH2AX) scores in a 1 Mb window around the break site calculated as the log2(gRNA1/tracrRNA only) for each replicate individually (dots) and all replicates combined (grey bar) in RPE-1 and K562 cells. (**D**) Correlation between ΔNL interaction and (ΔgH2AX scores, calculated as the log2(gRNA/tracrRNA only) ratio in RPE-1 (n = 7), and K562 (n = 2) cells. Each dot represents a 20 kb bin of a 15 Mb genomic region around the break site (chr7:80.2-95.2 Mb). Each graph shows the linear regression fit in blue and the Pearson correlation coefficient (R) in the top-left corner. (**E**) ΔNL interaction and ΔgH2AX scores, 10 (n = 7), 24 (n = 7), and 48 (n = 2) hours after gRNA1 transfection. The differential scores are calculated and visualized as those in Figure 2C.

Next, we tested whether the γH2AX domain matches the region that detaches from the NL. For this, we computed the differential γH2AX (ΔγH2AX) and NL interaction (ΔNL) scores across a 15 Mb region around LAD1 in consecutive windows of 20 kb and analyzed the correlation. Indeed, changes in NL interactions and accumulation of γH2AX changes showed a strong negative correlation (**Figure 3D**, Pearson’s R = –0.9). This analysis confirms that regions that lose NL interaction are marked by γH2AX. Additionally, we mapped γH2AX in K562 cells (**Figure 3B**) and observed similar correlations (**Figure 3D**, Pearson R = –0.74).

Next, we monitored whether the re-establishment of NL interactions coincided temporally with γH2AX clearance. Similarly to NL interaction re-attachments, γH2AX accumulation was partially reduced at 24 hours (**Figure 3E**) and there was no residual γH2AX measurable at 48 hours (**Figure S4D**). Thus, NL interaction dynamics correlate well with γH2AX dynamics in multiple cell lines.

### Tethering of DSBs to the NL slows down their repair and produces more unrepaired DSBs

We next investigated whether detachment from the NL is required for the repair of DSBs in LADs. Forced tethering of a genomic locus to the NL has been achieved previously^36^, but we aimed to force tethering to the NL immediately *after* DSB formation. To this end, we created a fusion protein consisting of Lamin B1 (LMNB1), green fluorescent protein (GFP) and the recently developed γH2AX-specific binding probe^48^ (_yH2AX_Reader-GFP-LMNB1). We hypothesized that this chimeric protein would be integrated in the NL and bind to DSBs that occur next to the NL, thus interfering with their detachment. To control for indirect effects of the binding of the γH2AX probe or of LMNB1 overexpression, also tested _yH2AX_Reader-GFP and GFP-LMNB1 fusion proteins (**Figure 4A**) We stably expressed each of the three constructs in RPE-1 cells and compared their subcellular localization by immunofluorescence to an RPE-1 cell line expressing GFP-NLS construct. As expected, the GFP-LMNB1 and _yH2AX_Reader-GFP-LMNB1 proteins were concentrated at the nuclear periphery, while the _yH2AX_Reader-GFP and GFP-NLS displayed a homogeneous nuclear localization (**Figure 4B & S5A**).

**Figure 4:**
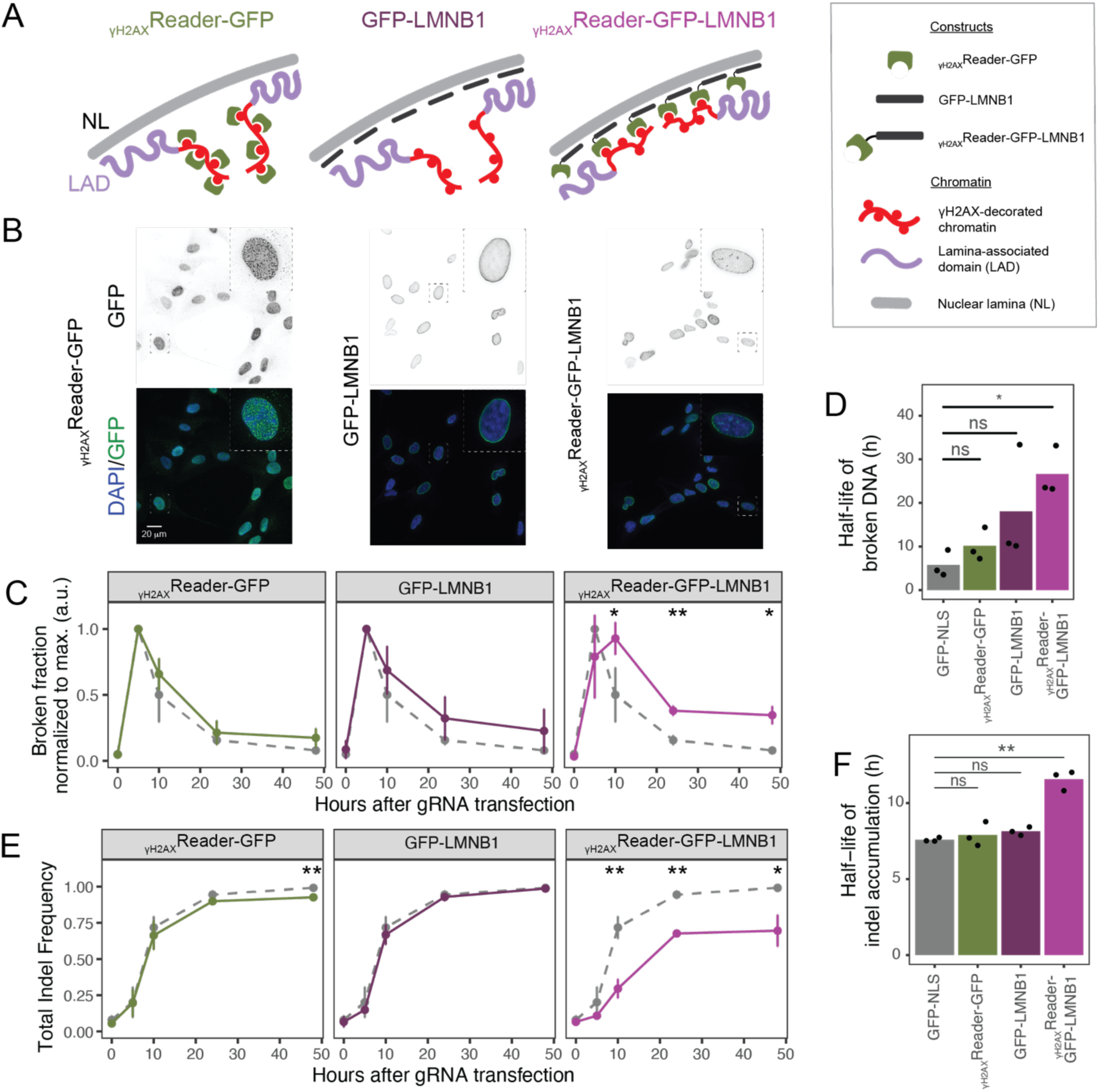
Tethering DSBs to the NL impairs repair. (**A**) Cartoon illustrating the effect of the NL tethering constructs. The γH2AX decorated chromatin around the DSB is depicted in red, the NL in grey and LADs in purple. From left to right: _yH2AX_Reader-GFP binds γH2AX decorated chromatin but does not interfere with the NL detachment (chromatin looping away from the NL). GFP-LMNB1 is localized at the nuclear periphery but does not interfere with the NL detachment because it does not bind γH2AX. _yH2AX_Reader-GFP-LMNB1 is localized at the nuclear periphery and binds γH2AX decorated chromatin next to the NL and prevents it from detaching. (**B**) Representative image of the GFP signal of RPE-1 cells overexpressing the NL tethering constructs. From top to bottom, GFP immunofluorescence signal and composed image of both channels (DAPI in blue and GFP in green). (**C**) Clearance of broken DNA measured by LM-PCR. The broken DNA measurements are normalized to the maximum broken fraction per timeseries. The measurements of the cell lines overexpressing the NL constructs are show in green and shades of purple as the mean of three biological replicates. The error bars represent the mean ± sd over three biological replicates. As reference in grey, the broken DNA measurements of a cell line expressing GFP-NLS construct is shown. The p-values of the differences in each timepoint between the reference and each of the constructs were tested using two-sided Student’s t-test. P-values that are lower than 0.05 are highlighted in the graph following the convention: “*” when p-val. < 0.05 and “**” when p-val. < 0.01. (**D**) The mean half-life of the broken 321 fraction calculated with a single-exponent decay function is shown as a bar-plot. Each dot shows the half-life of each biological replicate. The p-value is calculated using a two-sided Student’s t-test. (**E**) Accumulation of indels as measured by TIDE and plotted as described for Figure 4C. (**F**) Effect of NL tethering on the time required to reach the 50% of the maximum indel mutation accumulation (half-life) calculated with a sigmoidal function. Data were plotted as described for Figure 4D.

We then monitored DSB repair kinetics in these cell pools by measuring the fraction of broken DNA at several time points up to 48 hours. The maximum fraction of broken DNA was comparable across the various cell lines, indicating that Cas9-dependent break formation is unaffected by the expression of the different constructs (**Figure S5B**). To correct for subtle differences in break induction between biological replicates, we normalized the fraction of broken DNA to the maximum measured in each time series (**Figure S5B**). Interestingly, we find that repair in LAD1 is delayed in the cells expressing _yH2AX_Reader-GFP-LMNB1 as compared to reference cells. Importantly, expression of _yH2AX_Reader-GFP and GFP-LMNB1 did not significantly change the kinetics of repair (**Figure 4C**). Next, we estimated the half-life of the DNA breaks by means of a single-exponent decay model. Due to the low number of timepoints taken, the estimation of the half-life is very sensitive to outliers (see GFP-LMNB1, **Figure 4D**). Nonetheless, this analysis showed that half-life of a LAD-associated DNA break is longer in cells expressing _yH2AX_Reader-GFP-LMNB1, than in any of the other cell lines (**Figure 4D**).

Remarkably, cells expressing _yH2AX_Reader-GFP-LMNB1 contained a significantly larger amount of residual DNA breaks at 48 hours after transfection (**Figure 4C**, p-value = 0.002 and 0.02 respectively). Possibly, these remaining breaks cannot be repaired at all, because the fraction of broken DNA decreased minimally between the 24 and 48 hour timepoints. To exclude that the observed differences were due to an abnormal cell cycle arrest, we monitored the cell cycle distribution of the different cell lines following gRNA1 transfection. Cells accumulated in G1 phase at 24 and 48 hour after gRNA1 transfection, but this response was similar in all cell lines (**Figure S5C**). Taken together, these data indicate that repair of LAD-associated DSBs is impaired when the broken DNA remains tethered to the NL, producing long term complications in repair.

Next, we used TIDE ^43^ to investigate if tethering of the broken DNA to the NL affects the accumulation of indels. In line with the slower repair kinetics, we observed that cells expressing_yH2AX_Reader-GFP-LMNB1 accumulate indels more slowly than reference cells (**Figure 4E-F**): at 48 hours we detected a ∼40% reduction of indels (**Figure 4E**). It is worth noting that cells expressing _yH2AX_Reader-GFP also show a slight but significant reduction in the total indel percentage compared to control cells (**Figure 4E**). In summary, our data indicate that tethering of DSBs to the NL slows down repair kinetics of DSBs in LADs and leads to persistence of the DSB after 48 hours in a subset of cells. Thus, efficient repair of DSBs in LADs requires detachment of the surrounding DNA from the NL.

### γH2AX orchestrates rewiring of NL interactions

Next, we sought to investigate the molecular determinants of the NL detachment. We hypothesized that megabase-scale detachment from the NL is more likely to depend on chromatin changes that accumulate in large domains around DSBs, than kilobase-scale changes like resection or recruitment of DNA proteins to the broken ends^5^. Because the NL detachment strongly correlated with accumulation of γH2AX, we first tested whether NL detachment depends on γH2AX. For this, we established two H2AX knock-out (H2AX^KO^) RPE-1 clonal cell lines. Loss of H2AX in human and mouse cell lines leads to impaired binding of several DNA repair proteins, but no major DNA repair defects have been reported^12^. As a H2AX wild-type control, we isolated two H2AX wild-type RPE-1 clones (H2AX^WT^). After confirmation of the genotype of the clones (**Figure S6A**), we checked γH2AX induction by fluorescence microscopy. Indeed, H2AX^KO^ cells did not accumulate γH2AX upon 4 Gy irradiation (**Figure 5A & Figure S7A**). In line with previous observations ^13^, we also observed fewer 53BP1 foci in H2AX^KO^ compared to H2AX^WT^ cells (**Figure 5B & Figure S7B**), but no difference in the number of Rad51 foci (**Figure S6B & Figure S7C**).

**Figure 5:**
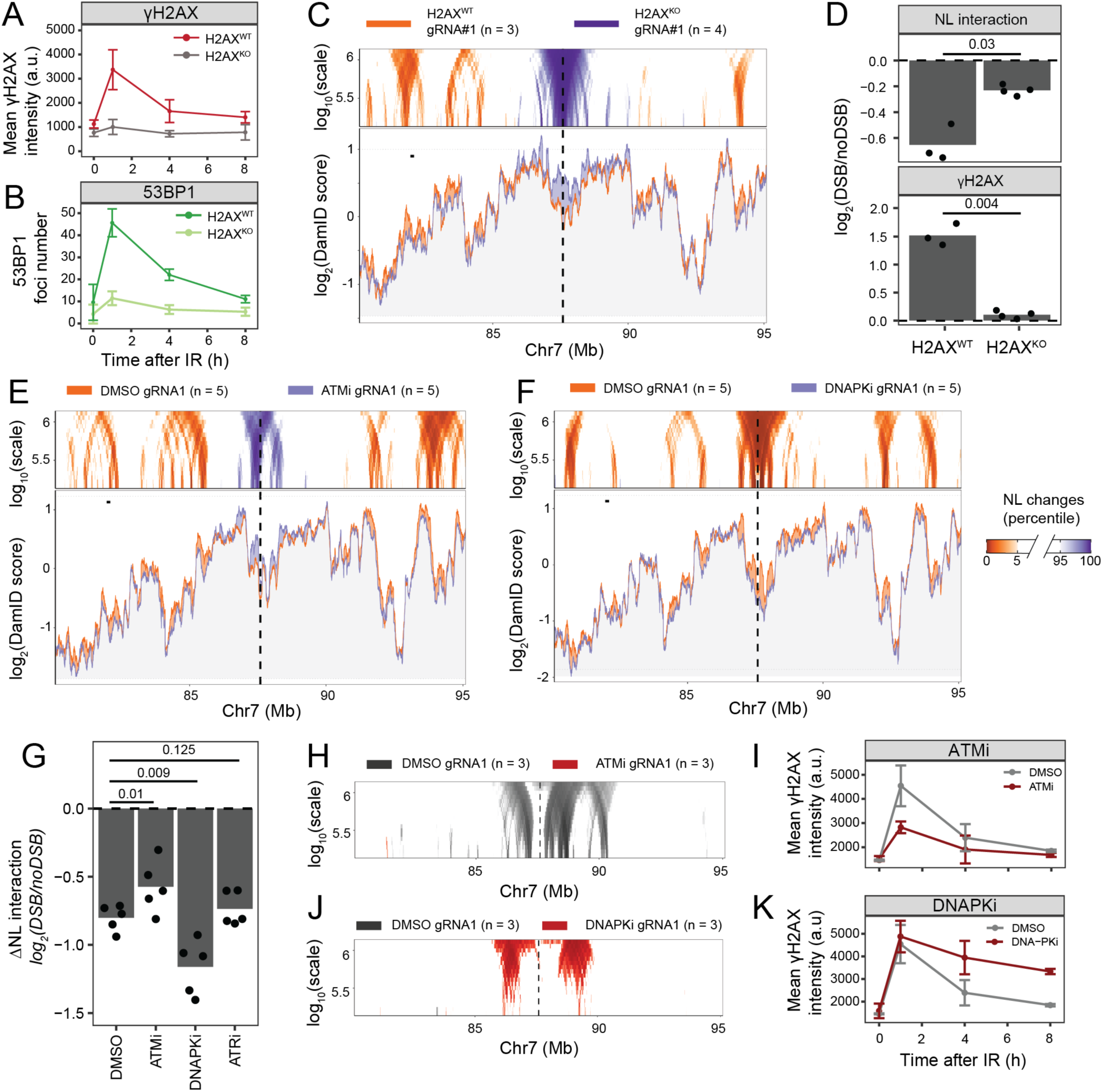
γH2AX orchestrates detachment from the NL. (**A**) Mean γH2AX nuclear intensity in H2AX^WT^and H2AX^KO^ RPE-1 cells at 1 hour (n = 6), 4 hours (n = 6), and 8 hours (n = 4) after 4 Gy irradiation or without irradiation (0 hours, n = 4). S phase cells were excluded from the analysis, based on EdU-positivity. Representative images are shown in **Figure S7A-B**. Error bars show the mean ± sd of the biological replicates. (**B**) Same as in Figure 5A but for the mean number of 53BP1 foci per cell. (**C**) Same as in Figure 1A, but for NL interaction differences between gRNA1 transfected H2AX^WT^ (orange, n = 3) and H2AX^KO^ (purple, n = 4) RPE-1 cells. In this graph, control and experimental samples are transfected with gRNA1. Therefore, the NL interaction changes represent the contribution of H2AX to NL detachment. (**D**) Quantification of the NL interaction (top) and γH2AX (bottom) differential scores, as calculated in Figure 1C, in H2AX^WT^(n = 3) and H2AX^KO^ (n = 4) RPE-1 cells. P-values are calculated using a two-sided Student’s t-test. (**E-F**) NL interaction changes of (**E**) ATM inhibitor (n = 5) and (**F**) DNAPK_cs_ inhibitor (n = 5) treated cells compared to vehicle-only treated RPE-1 cells. The changes are visualized as described in Figure 5C. (**G**) ΔNL interaction scores, as calculated in Figure 1C, in cells treated with ATM inhibitor, DNAPKcs inhibitor, ATR inhibitor compared to cells treated with vehicle-only (n = 5). p-values are calculated using a two-sided paired Student’s t-test between vehicle-only treated (reference group) and inhibitor-treated samples. (**H**) Domainogram visualization as described in Figure 3A of gH2AX changes in ATM inhibitor-treated cells compared to vehicle-only treated cells (n = 3). Representative images are shown in **Figure S7D**. (**I**) gH2AX kinetics after 4Gy IR in ATM inhibitor and vehicle-only treated cells. Same plot as in Figure 5A. (**J-K**) Same plots as in H and I, but for DNAPKcs inhibitor-treated cells.

Next, we tested whether NL detachment depends on H2AX. For this, we transfected gRNA1 and mapped NL interactions and γH2AX in H2AX^WT^ and H2AX^KO^ clones. After confirming that H2AX^KO^ clones do not accumulate γH2AX in LAD1 (**Figure S6C**) and excluding that loss of H2AX affects genome-wide NL interactions (**Figure S6D**, Pearson R = 0.89), we determined the degree of NL detachment. Visualization of NL interactions revealed that NL detachment is greatly reduced in H2AX^KO^ cells compared to H2AX^WT^ cells (**Figure 5C**). Although a slight NL detachment could be detected in H2AX^KO^ cells, (**Figure S6E**), quantitative analysis showed that the LAD remained significantly more attached in H2AX^KO^ cells (**Figure 5D**, p-value = 0.03). We conclude that NL detachment is largely dependent on the presence of γH2AX. Based on these results and the striking overlap of NL detachment with γH2AX accumulation (**Figure 3A**), we propose that H2AX-phosphorylation plays a crucial role.

### Detachment from NL partially depends on ATM and is antagonized by DNAPK_cs_

To further elucidate the role of γH2AX in the NL detachment, we sought to identify the kinase(s) involved in this process. γH2AX is primarily phosphorylated by the kinases ATM, DNAPK_cs_, and ATR during the DSB response^7–9^. Selective kinase activity inhibitors for these three proteins (KU5593, M3814, and VE-822, respectively) are available. Thus, we measured DSB-induced detachment of LAD1 from the NL in the presence of 10 µM KU5593, 1 µM M3814, 200 nM of VE-822 or vehicle-only. After ruling out that these inhibitors cause genome-wide changes in NL interactions themselves (**Figure S8A-C**), we tested the effects of each inhibitor on NL detachment 10 hours after gRNA transfection (**Figure 5E-F & S8D**). Inhibition of ATM reduced the detachment of LAD1 from the NL subtly but significantly (**Figure 5E**, p-value = 0.01), showing that kinase activity of ATM is partially responsible for NL detachment in RPE-1 cells. We note that the effect of ATM inhibition on NL detachment varied substantially between replicates (**Figure 5G**). In contrast, DNAPK_cs_ inhibition boosted NL detachment by ∼ 25% (**Figure 5F**, p-value = 5.26×10^−6^), indicating that DNAPK_cs_ antagonizes NL detachment. Lastly, ATR inhibition did not detectably affect detachment from the NL (**Figure S8D**) nor γH2AX accumulation (**Figure S8E**). Because this concentration of VE-822 is known to effectively inhibit ATR^49^, we conclude that ATR does not have a major impact on NL detachment. These data show that NL detachment is dependent on ATM and antagonized by DNAPK_cs_ activity.

Although both ATM and DNAPK_cs_ phosphorylate H2AX^9^, they affect NL detachment of LAD1 in opposite ways. To further explore the effect of these inhibitors on H2AX phosphorylation, we (i) mapped γH2AX by pA-DamID after gRNA1 transfection and (ii) measured γH2AX accumulation kinetics upon 4 Gy irradiation by immunofluorescence microscopy. Treatment with ATM inhibitor partially reduced γH2AX accumulation in LAD1 (**Figure 5H & S8F**) and global γH2AX accumulation after 4 Gy irradiation (**Figure 5I**). In contrast, DNAPK_cs_ inhibition led to increased γH2AX accumulation in LAD1 (**Figure 5J & S8G**). Interestingly, DNAPK_cs_ inhibition did not affect global γH2AX intensity one hour after irradiation but slowed down γH2AX clearance (**Figure 5K**). This persistence in γH2AX could explain the higher γH2AX accumulation in LAD1 and the effect on NL detachment in DNAPK_cs_-treated cells measured at 10 hours after break induction. These results are generally concordant with a model in which H2AX-phosphorylation induces NL detachment.

### Detachment from NL is not dependent on BRCA1

Previous studies have shown that resection, affecting kilobases of DNA adjacent to the break site^50^, contributes to increased mobility of DSBs in heterochromatin^28^. Although it is difficult to conceive how kilobases of resected DNA could induce megabase-scale rewiring of NL interactions, we tested whether NL detachment is dependent on resection. Because BRCA1 is a DNA repair protein that promotes resection^51,52^, we decided to map NL interactions after gRNA1 transfection in available RPE-1 p53^KO^ cells lacking BRCA1 (p53/BRCA1^KO^) and in RPE-1 p53^KO^ cells as controls (p53^KO^). Both cell lines showed a very similar NL detachment pattern as we observed in RPE-1 p53^WT^ cells (**Figure S9A-C**). Direct comparison only revealed subtle differences spread throughout the locus (**Figure S9D**). The minor difference in NL contacts that we observed between p53^KO^ and p53/BRCA1^KO^ cells after DSB formation were highly similar to the differences we observed between p53^KO^ and p53/BRCA1^dKO^ cells at baseline (see **Figure S9E**). This indicates that the modest BRCA1-associated changes that we observe are likely a consequence of the role of BRCA1 regulating heterochromatin rather than its role on DNA repair^53,54^. Furthermore, because recruitment of RAD51 was greatly reduced in cells lacking BRCA1 (**Figure S9F**), it is unlikely that resection is required for NL detachment.

### γH2AX accumulation at chromosome ends influences their NL association

Finally, we wondered whether γH2AX antagonizes NL interactions in other contexts than Cas9-induced DSBs. Previous studies have shown that γH2AX accumulates at telomeres and subtelomeric regions in several cell lines and model organisms^47^ ^Seo,2012,22467212,55,56^. These parts of the genome interact with the NL in early G1, but move towards the nuclear interior as cells progress through the cell cycle ^57^. We hypothesized that, similar to what we observe on DSBs, γH2AX could help detach chromosome ends from the NL. To study this, we first investigated whether distal parts of chromosomes are enriched for γH2AX in RPE-1 cells. For this, we applied a hidden Markov model to identify γH2AX domains in an unbiased manner. This analysis showed that all chromosome ends of non-acrocentric chromosomes accumulate γH2AX (95% confidence interval (CI) [0.4:7.1 Mb], **Figure 6A-B**). However, their signal was 50% weaker than the γH2AX domain measured in LAD1 after gRNA1 transfection (**Figure 6C**).

**Figure 6:**
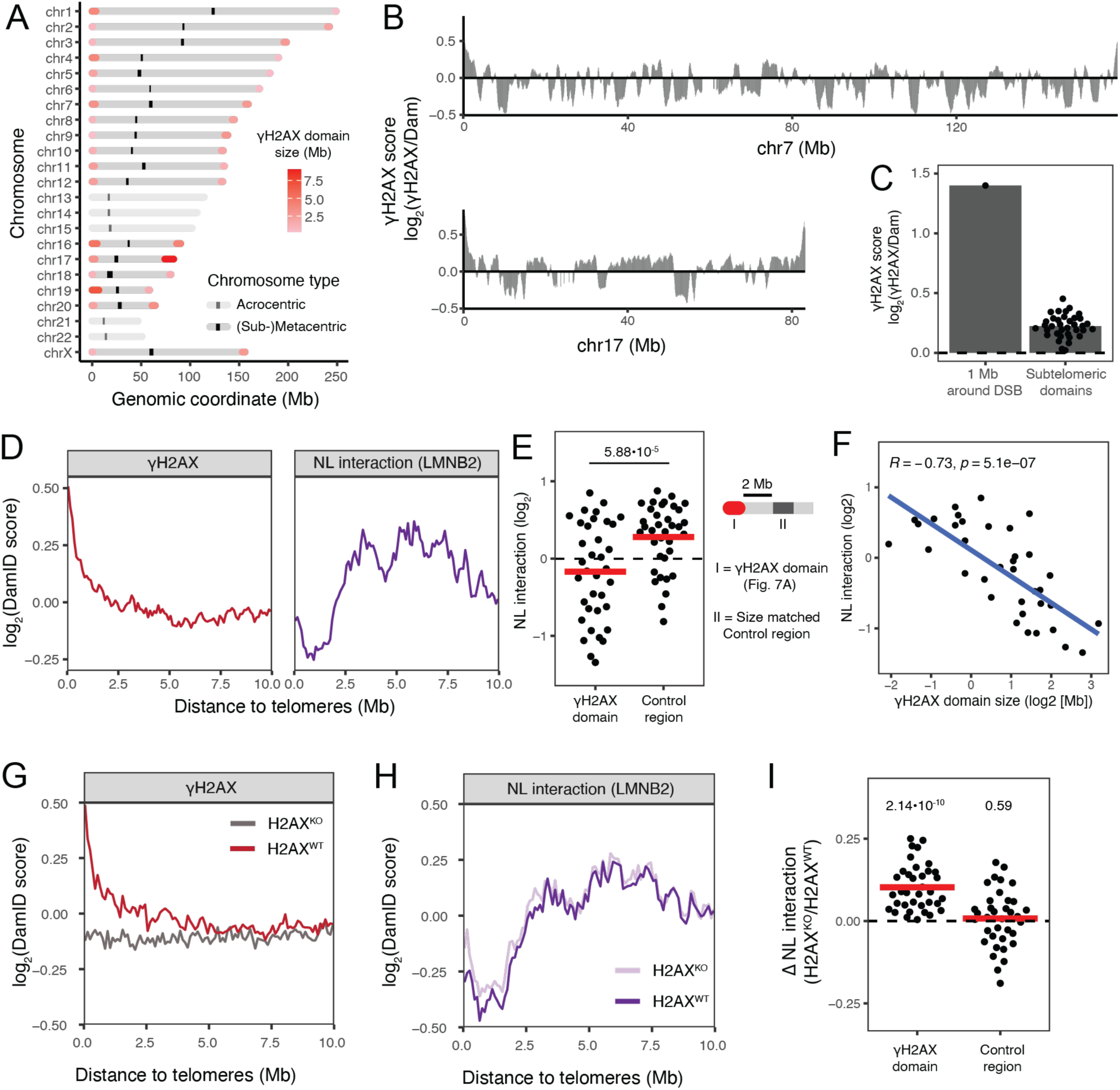
γH2AX accumulation at chromosome ends influences their NL association. (**A**) Genomic coordinates of the subtelomeric γH2AX domains at the ends of non-acrocentric chromosomes (solid colors) color in a scale of red reflecting their size in Mb. The centromeric repeats of each chromosome are plotted as black rectangles. (**B**) Whole chromosome log_2_(γH2AX/Dam) scores from tracrRNA only-transfected RPE-1 cells at 80 kb bin size resolution (n = 7). (**C**) Mean log_2_(γH2AX/Dam) values of subtelomeric γH2AX domains (variable size) in gRNA1 transfected cells. Each dot represents the mean value of a γH2AX domain in non-acrocentric chromosomes (n = 36) and the grey bar shows the mean of all domains. As a reference, the mean log_2_(γH2AX/Dam) score of a window of 1 Mb around the break is plotted. (**D**) Mean γH2AX (left) and NL (right) interaction scores (log_2_(DamID)) of chromosome ends in tracrRNA only-transfected RPE-1 cells. This plot shows the mean DamID scores as a function of distance to telomeres ([0-10] Mb) only from chromosome ends of non-acrocentric chromosomes. (**E**) Mean NL interaction scores (log_2_(LMNB2/Dam)) from each subtelomeric γH2AX domain (n = 36) in RPE-1 cells. The horizontal line represents the average NL score over all γH2AX domains. As a control for NL interactions of regions outside the γH2AX domain, we calculated the mean NL interaction score in a size-matched region 2 Mb away from the γH2AX domain, as illustrated in the cartoon on the right. The p-value was calculated using a two-sided paired Student’s t-test. (**F**) Correlation between mean NL interaction scores (log_2_(LMNB2/Dam)), as in Figure 6E, and subtelomeric gH2AX domain size in RPE-1 cells. The graph shows the linear regression 462 fit as a blue line, the Pearson correlation coefficient (R), and the p-value (p) of the linear regression in the top-left corner. The same plot as in Figure 6D, for (**G**) gH2AX scores and (**H**) NL interaction scores in H2AXWT (n = 3) and H2AXKO (n = 4) RPE-1 cells. (**I**) ΔNL interaction scores of subtelomeric gH2AX domains and adjacent control regions in RPE-1 cells (n = 36). The ΔNL interaction scores are calculated as NL interaction log2(ratio) between H2AXKO and H2AXWT cells. The horizontal red line represents the mean ΔNL interaction scores from each group and the dashed horizontal line shows no difference between H2AXKO and H2AXWT cells. p-values are calculated using a Student’s t-test where the null hypothesis is that ΔNL interaction scores are zero.

Next, we tested whether γH2AX antagonizes NL association at chromosome ends. For this, we first plotted the mean NL interaction and γH2AX scores over a 10 Mb window in the distal part of non-acrocentric chromosome ends (**Figure 6D**). This analysis revealed that subtelomeric regions decorated with γH2AX show low NL interactions that sharply increase outside the γH2AX domain. Quantitative analysis revealed that subtelomeric γH2AX domains have significantly lower NL interaction scores than adjacent control regions (**Figure 6E**, p-value = 5.58×10^−5^). Because of the large variability of NL scores of chromosome ends (95% CI [-1.27,0.74 log_2_]), we hypothesized that a critical mass of γH2AX could be needed to dissociate from the nuclear periphery. To test this hypothesis, we analyzed the correlation between the mean NL interaction scores (log_2_score) and the size of the γH2AX domain (log_2_[Mb]). This revealed a strong negative correlation between the NL interaction score and γH2AX domain size (**Figure 6F**, Pearson’s R = –0.73). We repeated these analyses for RPE-1 p53^KO^ and K562 cell data and found that γH2AX accumulation negatively correlates with the NL interaction of chromosome ends in these cell lines (**Figure S10,** p-value = 1.58×10^−4^ and 0.008, respectively). However, we note that the γH2AX domain size did not correlate with the NL interaction of telomeric regions in K562 cells (**Figure S10F**, p-value = 0.89). Nevertheless, we found that γH2AX accumulation negatively correlates with the NL association of chromosome ends in multiple cell lines and independently of the p53 status of the cell.

To directly test whether chromosome ends dissociate from the NL in a γH2AX-dependent manner, we analyzed their NL interaction in H2AX^KO^ clones, which lack the subtelomeric γH2AX domains (**Figure 6G**). Mapping of genome-NL interactions in H2AX^KO^ cells showed a subtle but significant increase of NL interaction scores at chromosome ends compared to H2AX^WT^ clones (**Figure 6H-I**, p-value = 2.14×10^−10^). In contrast, the NL association of adjacent regions remained unchanged in H2AX^KO^ clones. (**Figure 6H-I**, p-value = 0.59). These results indicate that γH2AX partially regulates the NL association of chromosome ends.

## DISCUSSION

Here, we report that DSBs in LADs lead to the transient NL detachment of megabases of the genome surrounding the break. Upon a Cas9-induced DSB, several megabases around the break site accumulate γH2AX, which appears to trigger NL detachment. However, detachment from the NL as not accompanied by loss of H3K9me2/3, indicating that the detachment does not depend on the removal of these heterochromatic marks. Moreover, we show that NL detachment is a widespread response to DSBs in LADs because it occurs across multiple LADs in the human genome and occurs independently of the p53 status of the cell. By tethering DSBs to the NL, we demonstrate that NL detachment is required for efficient repair, and failure to detach leads to an increase in the fraction of unrepaired DSBs. Finally, we show that γH2AX accumulation in sub-telomeric regions also counteracts NL interactions.

### A single DSB-induced loss NL detachment of megabases of the genome

Earlier studies showed that local detachment from the NL can be induced by chromatin de-compaction^58^ or gene activation^40,59^. Combined with the findings we report here, this implies that NL detachment is a common cellular response to different stimuli. However, the NL detachment caused by a single DSB spreads over ten-fold longer distances than the NL detachment induced by gene activation, which typicallyspreads over 50-100 kb. Moreover, we show that the DSB-induced NL detachment can spread to neighboring LADs. Another difference is that DSB-induced NL detachment recovers within 48 hours. This contrasts with the effect of chromatin decondensation, which can alter the nuclear positioning of a locus for at least 7 days^58^. Because of the temporal kinetics of DSB-induced NL detachment, we speculate that recovery of NL interaction requires the removal of γH2AX and re-establishing genome-NL lamina interactions after the subsequent mitosis^60^.

Previous studies have shown that DSBs in heterochromatin move to more repair-permissive chromatin contexts. These studies have described at least two types of movements: (i) movement to the edge of a microscopically defined heterochromatin domain^24,28,61–64^ and (ii) directed movement towards the nuclear periphery^27,32^. Here, we show that DSBs in LADs move an average of ∼ 250 nm towards the nuclear interior. Possibly, the DSBs move towards the edge of the LAD, which typically forms a chromatin layer of several hundred nm thick^60^. Although DSBs in a LAD cause a microscopically detectable shift towards the nuclear interior, we cannot exclude that a small fraction of DSBs move from the NL to a nearby nuclear pore^31^.

In addition to the NL detachment, we suggest that inter-LAD regions in the targeted locus gain NL interactions. These gains of NL interactions are scattered across several inter-LAD regions and in case of LAD1, they add up to 3.4 Mb. The amount of DNA that gains NL interaction is roughly similar to the amount that loses NL interactions, which suggests that these gains are “compensatory” movements caused by the vacated space at the NL. Similar “compensatory” movements have been detected after gene reactivation^40^, but these occurred at smaller scales. Moreover, ATM inhibition and H2AX KO, which both reduce the NL detachment, also reduced the NL interaction gains of inter-LADs. Thus, the NL interaction gains of inter-LAD regions correlate well with the NL detachment of LAD1.

### Failure to detach from the NL impairs DSB repair

By measuring the kinetics of DNA repair, both by determining the fraction of broken DNA and by determining accumulation of indels in cells expressing the NL tethering construct, we found that the detachment from the NL is required for efficient repair of DSBs. This suggests that LADs form a chromatin context that is relatively refractory to repair of DSBs. This could explain why DSBs in LADs are repaired slower and have pathway preferences that are different from other chromatin contexts^35,36^. Moreover, our experiments show that indel accumulation frequency is slower in cells in which we artificially tether the broken DNA to the NL. It is important to note that TIDE relies on a PCR reaction spanning the break site, causing broken DNA to remain undetected in TIDE assays. TIDE measures the fraction of indels relative to non-broken and/or error-free repaired sites. Therefore, we hypothesize that the lower indel accumulation could be caused by at least two phenomena: (i) a slowdown of all DNA repair pathways or (ii) a shift from mutagenic repair to non-mutagenic pathways. The latter would be in line with recent findings that suggest that DSBs from transcribed areas relocalize to the nuclear periphery to engage in HR^31^. If the latter hypothesis were true, it would imply that the movement away from the nuclear lamina would be associated to mutagenic DNA repair pathways such as NHEJ and/or MMEJ rather than HR.

### H2AX phosphorylation orchestrates detachment from NL

Our observations that H2AX knock-out and ATM kinase inhibitor treatment both reduce the detachment from the NL supports an active role of γH2AX. However, we cannot formally exclude that other H2AX modifications or H2AX itself induce detachment from the NL. While previous studies indicated that γH2AX accumulates in the interior of heterochromatic domains before relocalization^24,28^, they never established an active role of γH2AX in the exclusion of DSBs from heterochromatin.

Although inhibition of ATM partially reduces the NL detachment, it does not entirely prevent γH2AX accumulation. The partial effect of ATM inhibition suggests that other kinases phosphorylate H2AX in LADs and might be involved in regulating the detachment of LADs from the NL. Our data rule out a contribution of DNAPK_cs_ in this process, but it is possible that combining ATM and ATR-inhibitors would lead to a further reduction in NL detachment. Since ATM inhibition is only leading to a partial reduction in H2AX phosphorylation, we hypothesize that multiple kinases might need to be inhibited simultaneously to completely ablate H2AX phosphorylation in a broken LAD and thereby fully prevent NL detachment. Nevertheless, the concordance between γH2AX levels and NL detachment across all three inhibitor treatments is in line with our hypothesis that H2AX phosphorylation levels dictate NL detachment.

Currently, we can only speculate how γH2AX accumulation leads to dissociation from the NL. One possibility is that H2AX phosphorylation increases the net negative charge of chromatin around the break site. Such a change over several megabases might lead to chromatin de-compaction and detachment from the NL. An alternative hypothesis is that γH2AX acts as a binding platform for one or more other DNA repair proteins, which in turn trigger NL detachment. A possible candidate would be MDC1, which binds to chromatin in a γH2AX-dependent manner^10,11^ and contributes to the recruitment of DNA repair proteins. Additional studies are required to understand how γH2AX triggers NL detachment.

In addition to its role in DSB-induced NL detachment, we report that γH2AX accumulation in subtelomeric regions influences the NL association of these regions. Although the causes of γH2AX accumulation in subtelomeres are not fully understood, it has been proposed that it is caused by γH2AX spreading from occasionally uncapped telomeres^55^. Because our analysis is limited to the mappable regions of the genome, we cannot determine whether γH2AX accumulation causes detachment of telomeric repeats from the NL. Nevertheless, our data indicates that γH2AX domains can influence NL association of genomic loci in a Cas9 independent setting.

### Limitations of this study

Although our data indicates that γH2AX antagonizes NL interaction in sub-telomeric regions in a Cas9-independent setting, most of our experiments use Cas9 to induce DSB in LADs. Several studies have shown that Cas9-induced DSBs are repaired more slowly than other DSBs^44,65^ and Cas9 post-cleavage residence time on DNA might affect DNA repair^66,67^. Taking all the caveats of Cas9 together, we cannot exclude that only Cas9-induced DSBs lead to detachment from the NL.

Another limitation of this study is the potential underestimation of the NL detachment effect size. First, the induction of DSBs in our system is not fully penetrant. We estimated that the uncut fraction is around 40% by measuring the percentage of wild-type sequences by TIDE. The low frequency of Cas9 cutting is most likely a result of the heterochromatic nature of the locus we are targeting. Second, Cas9-dependent DSB induction is asynchronous. We estimated that only maximally 10% of the gRNA1 target sequences are broken at any given timepoint after gRNA1 transfection. As the pA-DamID maps report on the average NL contacts in a large cell population, it is very likely that we underestimate the degree of NL detachment.

Furthermore, we performed our experiments in a cell cycle agnostic manner. Thus, we do not know whether the NL detachment is limited to a specific cell cycle phase or occurs throughout the cell cycle. Because one study found that relocalization of DSB away from heterochromatin is limited to G2 phase^28^, this might also be the case for NL detachment.

We also did not explore the DNA repair pathways that act on the break following its detachment from the NL. Although ATM inhibition and H2AX knock-out favor MMEJ over NHEJ in Cas9-induced breaks^37^, the relation of these findings to the NL detachment we report here is not entirely clear. Moreover, the fact that NL detachment is not dependent on resection indicates that it must occur upstream of resection or be associated to NHEJ. Future studies may address these unresolved issues and provide more mechanistic insights into the detachment of entire LADs from the NL as the result of a single DSB.

## METHODS

### Cell culture

For this study, we primarily used the RPE-1 iCut cell line (RPE1) ^41^, which expresses Cas9 in a doxycycline– and Shield-1-inducible manner. Cells were cultured in DMEM/F12 (1:1) medium (GIBCO, cat. no. 11320-033) supplemented with 10% fetal bovine serum (FBS, Capricorn Scientific) and 1% penicillin/streptomycin (GIBCO, cat. no. 15070-063) at 37 °C and 5% CO2. Additionally, we used RPE-1 p53^KO^, RPE-1 p53/BRCA1^dKO68^ and K562 cells^35^. The RPE-1 p53^KO^ and p53/BRCA1^dKO^ cell lines are clonal knock-out cell lines that constitutively express Cas9, and were cultured using the same conditions as RPE-1 iCut cells. K562 cells, expressing Cas9 in a Shield-1-inducible manner were cultured in RPMI 1640 (GIBCO, cat. no. 1187-093) supplemented with 10% fetal bovine serum and 1% penicillin/streptomycin. We regularly tested the cells to be free of mycoplasma.

### pA-DamID experiments

#### gRNA transfection in RPE-1-derived cells

To harvest at least 0.5 million RPE-1 cells per pA-DamID sample, we optimized the number of cells seeded per cell line. Because of differences in growth rate, we plated 1.25 million RPE-1 and RPE-1 p53KO and 1.5 million RPE-1 p53/BRCA1dKO cells a day prior to lipofection. Cells were plated on 10 cm dishes in 8 mL of DMEM:F12 medium supplemented with 500 nM Shield-1 (Aoubious, cat. no. AOB1848) and 1 µM doxycycline (Sigma Aldrich, cat. no. D9891-1G) to induce Cas9 expression. The following day, we transfected gRNA1, gRNA2, and 15-nt gRNA1 (**see Table 1 for sequences**) at a final concentration of 20 nM and 1:500 dilution of RNAiMAX in an 8 mL final volume. For this, we combined 0.16 nmol of crRNA with 0.16 nmol of tracrRNA in 40 µL Duplex Buffer (IDT, cat. no. 1072570) and incubated the mix for 3 minutes at 95 °C. While incubating the crRNA:tracrRNA mix, we diluted the RNAiMAX lipofectamine (ThermoFisher, cat. no. 13778150) 1:50 in 800 µL of Optimem (GIBCO, cat. no. 31985070). Next, we incubated both mixes at room temperature for 5 minutes and mixed the crRNA:RNA and lipofectamine solutions. After incubating for 15 minutes at room temperature, we added the transfection mix dropwise to the cells. For the mock-transfected samples, we did not add the crRNA and transfected only the tracrRNA following the same protocol.

For the pooled gRNA experiments (pool 9), we combined the ten gRNAs and transfected them at a final concentration of 200 nM (20 nM of each gRNA) and 1:50 dilution of RNAiMAX in an 8 mL final volume. We used 10 times more gRNA, tracrRNA and RNAiMAX lipofectamine then previously used for single gRNA transfection. Otherwise, we followed the same protocol used for single gRNA transfections.

#### gRNA nucleofection in K562 cells

To transfect gRNA1 in K562 cells, we nucleofected Cas9 ribonucleoprotein (RNP) with the SF cell Line 4D-Nucleofector X Kit S (Lonza, cat. no. V4XC-2032) and a 4D-Nucleofector X (Lonza, cat. no. AAF-1003X). We performed three nucleofection reactions per experimental condition because each reaction is optimized for 300,000 cells, and we need at least 0.5 million cells to perform pA-DamID. For each individual nucleofection reaction, we mixed 0.12 nmol of gRNA1 crRNA and 0.12 nmol tracrRNA in 6.6 µL of Duplex Buffer and incubated at 95 °C for 5 minutes. Then, we added 0.1 nmol of purified Cas9 protein (IDT, cat. no. 1081059) and incubated the mix for 15 minutes at room temperature. Next, we combined 300,000 K562 cells, 0.1 nmol Cas9 RNP mix, and 1.3 µL of a 100 µM electroporation enhancer solution (IDT, cat. no. 1075915) in 20 µL SD cell line solution (Lonza, cat. no. PBC2-00675) in a 16-well nucleofection strip and electroporated with the FF-120 program. After incubating the nucleofected mix for 10 minutes at room temperature, we added 75 µL of pre-warmed (37°C) RPMI medium complete medium. Then, we combined three reactions per pA-DamID sample in a 6-well plate with 2 mL of RPMI complete medium supplemented with 500 nM Shield-1.

#### Inhibitor treatments

For the DNA repair kinase inhibitor pA-DamID experiments, we treated RPE-1 cells with a final concentration of either 10 µM ATM inhibitor (KU5593, from a 10 mM stock in DMSO, Calbiochem cat. no. #118500), 1 µM DNAPK_cs_ inhibitor (M3814, from a 1 mM stock in DMSO, MCE cat. no. HY-101570), 200 nM ATR inhibitor (VE-822, from a 200 µM stock in DMSO, Selleckchem cat. no. S7102) or 1:1000 DMSO-only vehicle control (Sigma cat no. D4540). We added one of the inhibitors or the DMSO-only vehicle control (Sigma Aldrich, cat. no. 472301) to the RPE-1 cells 30 minutes before gRNA transfection and kept the inhibitors in the media throughout the experiment.

#### H2AX^KO^ RPE-1 cell generation

We generated clonal H2AX knock-out cell lines in RPE-1 iCut cells using CRISPR/Cas9-mediated gene editing. We designed two gRNAs targeting the first exon of the H2AFX gene (**for sequences see Table 1**) and seeded 100,000 RPE-1 cells for each gRNA transfection with DMEM:F12 complete medium supplemented with 500 nM Shield-1 and 1 µM doxycycline to induce Cas9 expression. A day after induction of Cas9, we transfected each gRNA separately at a final concentration of 20 nM following the protocol described previously, but this time in a final volume of 2 mL. Three days after gRNA transfection, we re-plated the cells in limiting dilution (average of 0.5 cells/plate) in a 96-well plate using conditioned medium (a mix of 45% complete DMEM:F12 medium, 45% of conditioned DMEM:F12 medium from exponentially growing RPE-1 cells, supplemented with 10% FBS). After couple of weeks of clonal expansion, we split the clonal cell lines and genotyped them using TIDE, using primers spanning the first exon of H2AFX (**for sequences see Table 1**). Two clones with frameshifting mutation and no presence of wild-type allele were selected based on Sanger sequencing traces. To control for variability between clones, we also generated two H2AX^WT^ clonal cell lines. For this, we transfected RPE-1 cells with tracrRNA only and we established two clonal cell lines and confirmed that they do not exhibit any mutation in the H2AFX gene. To assess the effect of H2AX^KO^ on NL detachment, we performed pA-DamID and fluorescence microscopy with each clone separately and considered each of them a separate biological replicate.

#### Generation of the NL tethering construct expressing cell lines in RPE-1

We generated RPE-1-derived cell lines stably expressing the NL-tethering and control constructs using lentiviral transduction. For this, combined the different elements using Gibson assembly in the construct in a pLGW-V5-EcoDam backbone (Addgene #59210) digested by AgeI and NheI restriction enzymes. We amplified the inserts by PCR using the NEBNext PCR reaction mix from different plasmids. The yH2AX-binding domain was PCR-amplified from a plasmid kindly provided by the Tuncay lab, and the mouse LMNB1 ORF was amplified from a Dam-LMNB1 plasmid that is used for DamID ^69^. The final constructs were validated by whole plasmid sequencing (Eurofins). Next, we produced lentivirus from each construct by cotransfecting 6 µg of each of the constructs, combined with 1.5 µg pMD2G (Addgene #12259) and 4.5 µg psPAX2 (Addgene #12260) of the packaging plasmids in HEK293T cells. The medium was changed 6 hours after transfection and we harvested virus-containing supernatant at 24 and 48 hours post transfection. To obtain RPE-1 cells stably expressing each of the constructs, we transduced 250,000 RPE-1 cells with 250 µL of lentivirus-containing supernatant. After at least a week of cell expansion, we sorted 150,000 GFP-positive cells into a 6-well and expanded the polyclonal population for cryopreservation. Lastly, we confirmed stable expression of the constructs by assessing the subcellular localization of the GFP marker of each construct under a fluorescence microscope (see below).

#### Detection of broken DNA by ligation-mediated PCR

We measured the fraction of broken DNA at different timepoints (5, 10, 24 or 48 hours) after gRNA1 transfection by ligation-mediated PCR (LM-PCR) as described in ^44^ with the following modifications. First, we resuspended the cells in 350 µL lysis buffer (1% SDS solution in 10 mM Tris-HCl pH 8.2 and 2 mM EDTA solution supplemented with 4 µL of 10 µg/ml proteinase K (Bioline, cat. no. BIO-37084) and 4 µL of 10 µg/ml RNAseA (ThermoFisher, cat. no. EN0531)) and incubated them for 3 hours at 55C. Next, we performed genomic DNA (gDNA) extraction by adding 40 µL of 3 M NaOAc and 1000 µL of 96% EtOH supplemented with 1.3 µL of 15 mg/mL GlycoBlue (ThermoFisher, cat. no AM9515). After gDNA extraction, we used 200 ng of gDNA with the extension primer 1 (XV432) in a 20ul single cycle PCR and ligated the dsDamID adaptor (XV426 and XV430 primers) to the blunted ends by incubating the mix with 15 units T4 Ligase (Roche diagnostics, 10799009001 5 U/µL) in a total volume of 60 µL overnight at 16 °C and inactivated the enzyme by incubating at 80 °C for 20 mins. After ligation, we quantified the frequency of broken DNA fraction using the adapter ligation efficiency as a proxy. We did this using two alternative methods: quantification of the PCR product in an agarose gel (gel quantification) or quantitative PCR (qPCR). Because the gel quantification and qPCR measurements showed a very good correlation (**Figure S1D**), we considered them as technical replicates and averaged them for downstream analysis.

To measure broken fraction by the gel quantification method, we performed two independent 30 µL PCR reactions using MyTaq 2x Red Mix (GC-Biotech, BIO-25043) with 5 µL (∼20ng) of the ligation mix each with the reverse primer 2 (XV431) and two different forward primers; the adaptor specific primer (XV428) or primer3 (XV430). The PCR reaction with the reverse primer and the adaptor specific primer amplified the fraction of DNA where the dsDamID adaptor ligated, while the other reaction (primer2-primer3) controls for loading differences. Then, we loaded 10 µL of the 30 µL PCR on a 1.5% agarose gel and quantified band intensity by drawing rectangles of equal size around the bands observed in the gel using the Image Lab software from BioRad. Finally, we measure the frequency of broken DNA by normalizing the intensity measurement of the adaptor specific PCR reaction to the ‘loading’ control sample after DNA band size correction.

To measure broken fraction by the qPCR method, we quantified the adapter ligation frequency running the two PCR reactions described previously using a Thermo Scientific Quantstudio 5 thermocycler and the following conditions. We performed these PCR reactions in a final volume of 10 µL with 2.5 µL of a 1:10 dilution of the ligation reaction as input, 2.5 µL of each primer set at 800 nM and 5 µL Sensifast SYBR no Rox (Gc-Biotech, BIO-98020) as follows: one cycle of 3 minutes at 95 °C, 45 cycles of 5 seconds at 95 °C, 10 seconds at 60 °C and 10 seconds at 72 °C. After running the amplification phase, we ran a melting curve by running a cycle of 5 seconds at 95 °C and 20 seconds at 60 °C followed by heating the sample up to 95 °C at a rate of 0.075 °C/s. We performed the data analysis using the Applied Biosystems design&analysis 2.8.0 software.

#### TIDE analysis of editing efficiency

We monitored editing efficiency of each gRNA in all pA-DamID experiments by TIDE^43^. While collecting cells for pA-DamID, we re-plated 100,000 cells with DMEM:F12 supplemented with doxycycline and Shield-1. 48 hours after re-plating, we collected the cells by trypsinization and lysed them in 50 µL of DirectPCR lysis buffer (Viagen cat. no. 301-C) supplemented with 1 mg/ml proteinase K (Bioline, cat. no. BIO-37084). Cells were incubated in lysis buffer for 16 hours at 55 °C, followed by an inactivation step for 45 min at 85 °C. After cell lysis, we amplified DNA fragments containing the gRNA target sites using primers annealing at ∼ 250 bp from the break site (for gRNA1: AGM11 and AGM12, **see Table 1**). We performed the PCR using 2 µL of lysate with 10 pmol of each primer and in a final volume of 20 µL of MyTaq RedMix (Bioline, cat. no. BIO25048) with the following conditions: 1 cycle at 95 °C, 30 cycles of 30 seconds at 95 °C, 15 seconds at 58°C and 30 seconds at 72 °C and 1 cycle of 1 min at 72 °C. Next, we degraded the excess primers by treating 10 µL of the PCR reaction with 0.125 mU of Shrimp Alkaline Phosphatase (New England Biolabs, cat. no. M0371S) and 0.25 mU of Exonuclease-I (New England Biolabs, cat. no. M0293S) in a final volume of 12.5 µL for 37 °C for 30 min followed by heat inactivation for 10 minutes at 95°C. Next, we Sanger sequenced 2.5 µL of the cleaned-up PCR product with 25 pmols of AGM11 in a final volume of 15 µL in ddH2O by Eurofins genomics. Lastly, we analyzed the resulting Sanger sequence traces with the TIDE algorithm to determine the editing frequency.

#### pA-DamID

pA-DamID was performed as described previously^38^ at different time points (10, 24, or 48 hours) after gRNA transfection in multiple RPE-1-derived cell lines and K562 cells. In brief, we harvested 0.5 to 1 million RPE-1 cells and 300,000 K562 cells per pA-DamID sample (antibody or Dam-only). After a first wash with PBS and a second one with digitonin wash buffer (20 mM HEPES-KOH pH 7.5, 150 mM NaCl, 0.5 mM, spermidine, 0.02% digitonin, – EDTA cOmplete Protease Inhibitor Cocktail), we incubated the cells with either anti-LMNB2 (Abcam, cat. no. ab8983), anti-γH2AX (Merck Milipore, cat. no. JDW301), anti-H3K9me2 (Acitve Motiv, cat. no. 39239), anti-H3K9me3 (Diagenode, C15410193) or anti-H3K27me3 (Diagenode, C15410195) antibodies at a 1:100 dilution in 200 µL digitonin wash buffer for two hours at 4°C. Unless stated differently, we performed every incubation and washing step with digitonin wash buffer. After two hours of incubation, we washed the cells and incubated them with anti-rabbit mouse antibody (Abcam, ab6709) at a 1:100 dilution in 200 µL wash buffer for one hour at 4 °C. Next, we washed the cells and incubated them with 200 µL of pA-Dam solution diluted 1:200 for 1 hour at 4 °C. After two washes, we resuspended the cells in 100 µL wash buffer supplemented with 80 µM SAM (New England Biolabs, cat. no. B9003S) and we activated the dam enzyme by incubating the cells at 37 °C for 30 minutes. To control for DNA accessibility and amplification biases of the Dam enzyme, we processed another sample with buffer only and incubated it with 4 units of Dam enzyme (New England Biolabs, M0222L) during the dam activation step. After a last wash, we pelleted the cells and stored them at –20 °C for downstream processing.

#### pA-DamID sequencing library preparation and sequencing data processing

To prepare the pA-DamID sequencing library, we first isolated gDNA from RPE-1 cells using the ISOLATE II genomic DNA kit (Bioline, BIO-52067) and eluted the genomic DNA in a final volume of 30 µL of ddH2O. Because we processed a low number of K562 cells per sample, we isolated the genomic DNA from K562 cells by ethanol precipitation. In brief, we lysed the cells with 530 µL lysis buffer (10 mM Trish-HCl pH 8.2, 400 mM NaCl, 2 mM EDTA, and 0.55% SDS) supplemented with 5 µL of 10 mg/ml proteinase K and 5 µL of 10 mg/ml RNAseA by overnight incubation at 55°C. After the lysis step, we added NaOAc to a final concentration of 300 mM, GlycoBlue (ThermoFisher, cat. no. 01216232) to a final concentration of 0.05 mg/ml, and 2.5 volumes of 96% ethanol. Next, we precipitated the genomic DNA by incubating the mix at –20 °C overnight followed by centrifugation at 30 min at 13,000 g. Subsequently, the pellets of genomic DNA were washed once with 70% ethanol and resuspended in 30 µL of ddH2O. After gDNA extraction, we processed up to 500 ng of gDNA for DNA sequencing. Because we used both Illumina HiSeq and Illumina NOVAseq platforms, we used different library preparation protocols depending on the DNA sequencer used in each experiment. The DNA sequencer and library preparation protocol used for each sample is described in GEO.

For samples sequenced in the Illumina HiSeq 5000 platform, we performed the sequencing library preparation and data processing as previously described^38^. In brief, we digested the genomic DNA with DpnI (New England Biolabs, cat. no. R0176L). Next, we ligated the DpnI adapter on the digested DNA and performed the Methyl PCR using DamID PCR primers. After the Methyl PCR step, we pooled the samples based on DNA quantity and purified them. Next, we blunted the 3’ DNA ends with the DNA end repair kit (Lucigen, cat. no. ER81050) and A-tailed the DNA with a Klenow polymerase fragment lacking exonuclease activity (New England Biolabs, M0212M). Then, we ligated the Y-shape Illumina adapter and added the Illumina indexes by PCR amplification. After sample pooling and DNA purification, we sequenced them on a HiSeq 5000 platform for 100-bp single-end reads. Lastly, we aligned and processed the sequencing data using the previously published computational pipeline (https://github.com/vansteensellab/Lamina_CellCycleDynamics/tree/master/Sequencing/damid/processing_pipeline/bin).

We performed the library preparation for samples sequenced in the Illumina NOVAseq 6000 platform as follows. First, we digested up to 500 ng of genomic DNA with 10 U of DpnI (New England Biolabs, cat. no. R0176L) in 10 µL of 1x CutSmart Buffer for 8 hours at 37 °C followed by a heat inactivation step at 80 °C for 20 minutes. Next, we A-tailed the digested DNA fragments (10 µL) with 12.5 U Klenow fragment lacking exonuclease activity (New England Biolabs, M0212M) in a final volume of 15 µL of 1x CutSmart supplemented with 0.33 µM dATP (ThermoFisher, cat. no. R0141). We ran the A-tailing reaction for 30 minutes at 37 °C and heat-inactivated the enzyme by incubating 20 minutes at 75 °C. Then, we ligated x-Gene Stubby Adapters (IDT) to the A-tailed DNA (15 µL) in a final volume of 30 µL. For this, we added 3 µL 10x T4 Ligase Buffer, 2.5 U of T4 DNA ligase (Roche, 10799009001), 12.5 pmols of x-Gene Stubby Adapters, and ddH2O up to 30 µL. We incubated the ligation mix for 16 hours at 16 °C followed by a heat inactivation step for 10 minutes at 65 °C. Next, we performed the Methyl-Indexed PCR on 4 µL of ligated DNA with 5 pmols of a unique pair of x-Gen Dual combinatorial Indexes (IDT) for each sample. We ran the Methyl-Indexed PCR reaction in a final volume of 40 µL of 1x MyTaq RedMix (Bioline, BIO-25048) as follows: one cycle of 1 minute at 94 °C, 15 cycles of 30 seconds at 94 °C, 30 seconds at 58 °C and 30 seconds at 72 °C and one cycle of 2 minutes at 72 °C. We purified the amplified DNA using cleanPCR (CleanNA cat. no. CPCR-0050) beads at 1.6:1 beads:sample ratio and sequenced on a NovaSeq 6000 platform with 100-bp single-end reads. Lastly, we aligned and processed the DNA sequencing data using the previously published computational pipeline (https://github.com/vansteensellab/pADamID_ABCB1_Novaseq).

### NL detachment visualization by fluorescence microscopy

We visualized detachment from the NL by measuring the shortest distance between the ANCHOR-sequence integrated into the LAD1 locus and the NL. For this experiment, we used an engineered RPE-1 cell line (RPE-1 iCut ANCHOR3) ^70^ in which the ANCH3 sequence was integrated 200 kb away from gRNA1 target site (chr7:87492431-87492874) and cells were engineered to express the OR3-GFP protein fusion. Because the OR3-GFP protein oligomerizes around the ANCH3 sequence, we assessed the nuclear positioning of LAD1 by measuring the shortest distance between the OR3-GFP focus and the NL. We performed this experiment in four biological replicates.

#### gRNA1 transfection of RPE-1 iCut ANCHOR3 cell line

To assess the nuclear positioning of LAD1 after gRNA1 transfection, we plated 250,000 RPE-1 iCut ANCHOR3 cells on 13 mm diameter coverslips (two per well of a 6-well plate) with medium supplemented with 500 nM Shield-1 and 1 µM doxycycline to activate Cas9 expression. A day after seeding the cells, we transected gRNA1 at a final concentration of 20 nM in a final volume of 2 mL as described above. Ten hours after gRNA transfection, we fixed and permeabilized the cells with 4% formaldehyde (Merck, cat. no. 104003) and 0.5% Triton X-100 solution (Sigma, cat. no. 93443) in PBS for 15 minutes at room temperature. As a control, we followed the same protocol, but we transfected tracrRNA only.

#### Immunofluorescence of RPE-1 iCut ANCHOR3 cell line

After fixation, we washed the cells three times with 0.5% Tween solution in PBS (PBS-T) and blocked them with 4% bovine serum albumin in PBS-T (blocking buffer) for 1 hour at room temperature. Next, we incubated the cells overnight at 4 °C with a 1:500 dilution of the Lamin B1 antibody (Abcam, cat. no. ab16048) in a blocking buffer. After three washes with PBS-T, we incubated the cells with a 1:600 dilution of anti-rabbit Alexa 647 secondary antibody (Thermo Fisher, cat. no. A21245) in a blocking buffer for 2 hours at room temperature. Next, we washed the cells three times with PBS-T and incubated them with 1 µg/ml of 4′,6-diamidino-2-fenilindol (DAPI) in PBS-T for 5 minutes at room temperature. Lastly, we dehydrated the coverslips with 70% ethanol and mounted them on glass slides with Prolong Gold (ThermoFisher, cat. no. P36934).

#### Image acquisition and calculation of the shortest distance to the NL

We acquired the images and measured the shortest distance between the OR3-GFP focus and the NL as described previously^70^. In brief, we imaged the cells with an LSM 980 Airyscan 2 on a Zeiss Axio Observer 7 SP inverted microscope. Next, we analyzed the images with the IMARIS analysis software (Oxford Instruments). In brief, we segmented the NL surface in 3D, detected the OR3-GFP focus in each cell, and extracted the shortest distance between the NL surface and the OR3-GFP focus.

### Measurements of repair kinetics after γ-irradiation

To assess the recruitment kinetics of DNA repair proteins to γ-irradiation-induced DSB, we quantified the mean γH2AX nuclear intensity and the number of 53BP1 and RAD51 foci at different time points after 4 Gy irradiation (1, 4 and 8 hours) or without irradiation (0 hours). We performed this experiment in multiple RPE-1-derived cell lines (two H2AX^KO^ clones, two H2AX^WT^ clones, p53^KO^ and p53/BRCA1^dKO^ cells) or RPE-1 cells treated with ATM inhibitor, DNAPK_cs_ inhibitor or vehicle-only (DMSO). For the latter experiments, we added one of the inhibitors or DMSO to the cells one hour prior to γ-irradiation at the same doses used for the pA-DamID experiment (see above), and we kept the inhibitors in the media throughout the experiment.

#### Gamma-irradiation of RPE-1 derived cells

For this experiment, we plated 250,000 cells on 10-mm glass coverslips. The next day, we irradiated the cells with a dose of 4 Gy using a Gammacell Exactor (Theratronics) with a ^137^Cs source. To mark cells in S-phase, we incubated the cells with 10 µM EdU (Thermo-Fisher, cat. no. A10044) solution for 15 minutes before fixation. Next, we fixed and permeabilized cells with 4% formaldehyde (Merk, cat. no. 1040021000) and 0.5% Triton X-100 (Sigma-Aldrich, cat. no. X100-100ML) solution in PBS for 10 min.

#### Subcellular localization of NL tethering constructs

For this experiment, we plated 250,000 cells on 10-mm glass coverslips. The next day we fixed cells with 2% paraformaldehyde (Electron Microscopy Sciences, cat. no. 15714) in PBS for 5 min. After three washes of 5 min with PBS, we permeabilized cells with 0.5% Triton X-100 (Sigma-Aldrich, cat. no. X100-100ML) in PBS for 5 min on ice and then 10 min at room temperature. Next, we washed the cells three times with PBS for 5 min.

#### Immunofluorescence protocol

We followed the immunofluorescence protocol described above with the following modifications in these experiments. First, we blocked the cells with a 5% BSA (Sigma, cat. no. A4503) solution and incubated the secondary antibody and DAPI solutions for one hour. We used 1:1000 mouse anti-γH2AX (Merk, cat. no. 05-636), rabbit anti-53BP1 (Novus Biologicals, cat. no. NB100-305), rabbit anti-RAD51 (Abcam, cat. no. ab63801), rabbit anti-GFP (Abcam, cat. no. ab290) primary antibodies solutions and 1:1000 anti-mouse Alexa 488 (Invitrogen, cat. no. A-11001), anti-mouse Alexa 568 (Invitrogen, cat. no. A-11004), anti-rabbit Alexa 488 (Invitrogen, cat. no. A-11008) anti-rabbit Alexa 568 secondary (Invitrogen, cat. no. A-11011) antibody solutions. Before mounting the coverslips, we incubated the cells with 1:1000 Alexa Fluor 647 azide (ThermoFisher, cat. no. A10277) and 100 mM ascorbic acid solution in the EdU staining buffer (100 mM Tris-HCl pH 8.5 and 1mM CuSO_4_) for 30 minutes at room temperature to stain the EdU-positive cells. After mounting the coverslips, we stored them at 4 °C until image acquisition.

#### Image acquisition and imaging data analysis

We imaged the coverslips with a THUNDER Imager equipped with a 63×/1.40–0.60 OIL Obj. HC PL APO objective and a deep-cooled 4.2 MP sCMOS camera (Leica Microsystems), and we collected ten z-planes with a 0.75 μm step. Next, we analyzed the acquired images using Fiji/ImageJ software^71^, and measured the nuclear intensity and foci number of γH2AX, 53BP1, and RAD51 in these images with a custom-built macro in Fiji/ImageJ that enabled automatic and objective foci analysis (https://github.com/BioImaging-NKI/Foci-analyzer). In brief, the *Foci-analyzer* macro segmented cell nuclei based on DAPI and measured the nuclear intensity and number of foci per nucleus for each fluorescent marker^41^. To exclude foci generated as a consequence of DNA replication, S-phase cells were excluded from the analysis by filtering out cells with an EdU intensity value above the 55^th^ percentile. We classified a cell as a G1 cell when its DAPI intensity was below the 25^th^ percentile, and we classified a cell as a G2 cell when its DAPI intensity was above the 75^th^ percentile. Lastly, we measured γH2AX and 53BP1 kinetics in G1 and G2 cells, but we only measured RAD51 foci number in G2 because they mainly accumulate in this cell cycle phase.

#### Cell cycle profile of NL tethering constructs

To assess the cell cycle distribution of the cells expressing the NL-tethering and control constructs following gRNA1 transfection, we plated 250,000 cells in a 6-well plate with medium supplemented with 500 nM Shield-1 and 1 µM doxycycline to activate Cas9 expression. A day after seeding the cells, we transected gRNA1 at a final concentration of 20 nM in a final volume of 2 mL as described above. To mark cells in S-phase, we incubated the cells with 10 µM EdU (Thermo-Fisher, cat. no. A10044) solution for 15 minutes before fixation. Five, ten, twenty-four and forty-eight hours after gRNA transfection, we fixed the cells with 4% formaldehyde (Merck, cat. no. 1040021000) in PBS for 10 minutes at room temperature. Next, we permeabilized cells with 0.5% Triton X-100 (Sigma-Aldrich, cat. no. X100-100ML) solution in PBS for 10 min. We subsequently incubated the permeabilized cells with a 5% BSA (Sigma, cat. no. A4503) solution and then we incubated the cells with DAPI and 1:1000 Alexa Fluor 647 azide (ThermoFisher, cat. no. A10277) and 100 mM ascorbic acid solution in the EdU staining buffer (100 mM Tris-HCl pH 8.5 and 1mM CuSO_4_) for 30 minutes at room temperature. Cell cycle analysis was carried out on CytoFLEX S (Beckman Coulter). A total of 10,000 events were acquired for each sample. Cell cycle distribution was calculated after appropriate gating of cell populations based on DAPI and EdU fluorescence intensity using FlowJo (BD Biosciences).

### Data visualization, computational and statistical analysis

#### Publicly available heterochromatin tracks and LAD calling in RPE-1 cells

To characterize the heterochromatic status of LAD1 and the adjacent regions, we used publicly available NL interaction, H3K9me3, H3K27me3, and replication timing data in RPE-1 cells. We downloaded the processed H3K9me3 and H3K27me3 pA-DamID data from GEO (GEO_ID: GSE248980) ^72^, the NL interaction (LMNB1) DamID (4DNES7QSJV2E, 4DNESHGTQ73M) ^38^ and replication timing (4DNFI9R1MQ7U, 4DNFIRNKABQJ, 4DNFIK2TA4DS and 4DNFIBKO9LI7) ^73^ data from 4D Nucleome data repository (https://data.4dnucleome.org). Then, we visualized DamID and pA-DamID data at a resolution of 20 kb as log2(antibody/Dam) tracks and the replication timing data at 5 kb resolution as log2(Late/Early) tracks. Moreover, we determined the LAD coordinates in RPE1(4DNES7QSJV2E, 4DNESHGTQ73M) and K562(4DNES6SM8UGO,4DNESTAJJM3X) ^74^ data at 80 kb bin resolution using hidden Markov modeling (https://github.com/gui11aume/HMMt) on the average NL interaction data for each cell line with data from^38^.

#### Domainogram visualization of pA-DamiD data

We visualized the NL interaction changes in a 15 Mb window around LAD1 (chr7:80.1-95.1 Mb) in two different ways. First, we visualized the control (orange) and experimental (purple) log_2_(LMNB2/Dam) data and highlighted the regions that gained NL interactions in purple and lost NL interaction in orange. To correct systematic biases between experimental and control data^40^, we applied a loess fit (span = 0.5) to the experimental and control data comparison, and we used this fit to correct the experimental data. A vertical dashed line was plotted to indicate the gRNA target site. Second, we visualized the data as a domainogram running a previously described pipeline ^40^. In brief, the domainogram pipeline analyzes changes between two pA-DamID GATC tracks at different bin sizes and highlights the bottom and top 5^th^ percentiles in orange and purple scales, respectively. Furthermore, we visualized changes in H3K9me3, H3K9me2, and H3K27me3 histone modification using the same approach, but we used different colors to visualize changes for each antibody. To visualize the changes in γH2AX intensity, we modified the domainogram pipeline as follows. First, we did not apply the loess correction to the data. Because the γH2AX pA-DamID scores around the break site are much higher than the pA-DamID scores in the rest of the genome, the experimental data correction would require the extrapolation of the loess fit. Thus, we did not apply the loess correction to avoid inaccurate data correction. Second, we only visualized the bottom and top 1^st^ percentiles in the domainogram visualization. We decided to apply a more stringent cut-off for γH2AX because the visualization of the top 5^th^ percentile vastly overestimated the specific γH2AX signal, which we measured by comparing H2AX^WT^ and H2AX^KO^ clones (**Figure S6C**). We must note that even this stringent cut-off might overestimate the γH2AX changes because we identified a domain that is three-fold larger than the specific γH2AX signal (**Figure S4B**).

#### Differential NL interaction and γH2AX quantification

To quantify the effect size of the NL detachment and γH2AX accumulation, we computed a differential pA-DamID score. First, we smoothed the 20 kb binned pA-DamID control and experimental tracks using the “runmean” function from *the caTools* R package and a moving window size of 9 bins (k = 9). Next, we calculated the differential score for each bin by subtracting the control log_2_(antibody/Dam) score from the experimental log_2_(antibody/Dam) score for each biological replicate separately. Lastly, we averaged the data in two ways: (i) we averaged the differential scores of a 1 Mb window around the DSB to assess the biological reproducibility of the NL interaction and γH2AX changes, and (ii) we averaged the biological replicates to visualize the differential NL interaction scores across LAD1.

#### Identification of subtelomeric γH2AX domains and adjacent control regions

We identified the genomic coordinates of the subtelomeric γH2AX domains of non-acrocentric chromosomes using hidden Markov modeling (https://github.com/gui11aume/HMMt). We first applied the model on the 20 kb binned γH2AX tracks of mock-transfected RPE-1 cells. Next, we selected the γH2AX domains adjacent to the telomeres of non-acrocentric chromosomes and named them subtelomeric γH2AX domains. We defined the start of the telomeric repeats of each chromosome as the first or last base pair of GRCh38 genome assembly and filtered out acrocentric chromosomes 13,14,15,21 and 22. Lastly, we quantified the mean NL interaction score and the mean γH2AX score of the subtelomeric γH2AX domains for each chromosome end. As a control region for the NL interaction score of chromosome ends, we defined a control region for each subtelomeric γH2AX domain. For this, we selected a size-matched region 2 Mb away from the end of the γH2AX domain, and we calculated the mean NL interaction scores in these adjacent control regions.

#### UMAP visualization

We visualized genomewide NL interaction differences between experimental conditions as a Uniform Manifold Approximation and Projection (UMAP) plot. We calculated the UMAP with the *umap* R package, and we plotted two UMAP dimensions as a scatterplot.

#### Ideogram visualization

We visualized pool 9 gRNA target sites and subtelomeric γH2AX domains as ideograms. For this, we extracted the chromosome sizes and the genomic coordinates of the centromeric repeats from the “human_karyotype” dataset of the *RIdeogram* R package^75^, which uses the GRCh38 genome assembly.

#### Cumulative plots

We visualized the average NL interaction and γH2AX scores from subtelomeric regions of non-acrocentric chromosomes as cumulative plots. For this, we calculated the distance of each bin to the telomere as defined above and averaged the NL interaction and γH2AX scores next by their distance to the telomere. Lastly, we visualized the mean score for the 10 Mb closest to the chromosome ends.

## DATA AVAILABILITY STATEMENT

All raw data shown in this paper have been deposited in GEO (BioProject no. PRJNA1274201, https://www.ncbi.nlm.nih.gov/bioproject/PRJNA1274201). Processed data is available in GEO and/or can be found on GitHub (https://github.com/vansteensellab/DSB_NL_detachment/import). GEO or 4DN accession numbers for publicly available datasets used in this manuscript are indicated in the methods section.

## CODE AVAILABILITY

All code to analyze the data and create the figures is available on GitHub (https://github.com/vansteensellab/DSB_NL_detachment/import).

## AUTHOR CONTRIBUTIONS

X.V., A.G.M., M.N.C, B.v.S and R.H.M conceived and designed the study. X.V., A.G.M, M.N.C, M.d.H, A.O., and S.G.M. performed experiments. X.V, M.N.C and T.v.S performed data analysis. R.C.S and, T.B. provided key reagents. B.v.S and R.H.M supervised the study. X.V., B.v.S and R.H.M. wrote the manuscript with input of all authors.

## ACKNOWLEDGMENTS

We thank the NKI Genomics and Research High Performance Computing core facilities for technical support; members of our laboratories for inspiring discussions and helpful comments. This work was supported by ZonMW TOP grant 91215067 (to R.H.M. and B.v.S.), European Research Council (ERC) Advanced Grant 694466 (to B.v.S); NWO Zwaartekracht (to R.H.M.); MSCA-IF 838555, and Next generation EU-MUR MSCA Young Researcher (to S.G.M). The Oncode Institute is partly supported by the Dutch Cancer Society. Research at the Netherlands Cancer Institute is supported by an institutional grant of Dutch Cancer Society and of the Dutch Ministry of Health, Welfare and Sport.

## SUPPLEMENTARY FIGURES

**Supplementary Figure 1:**
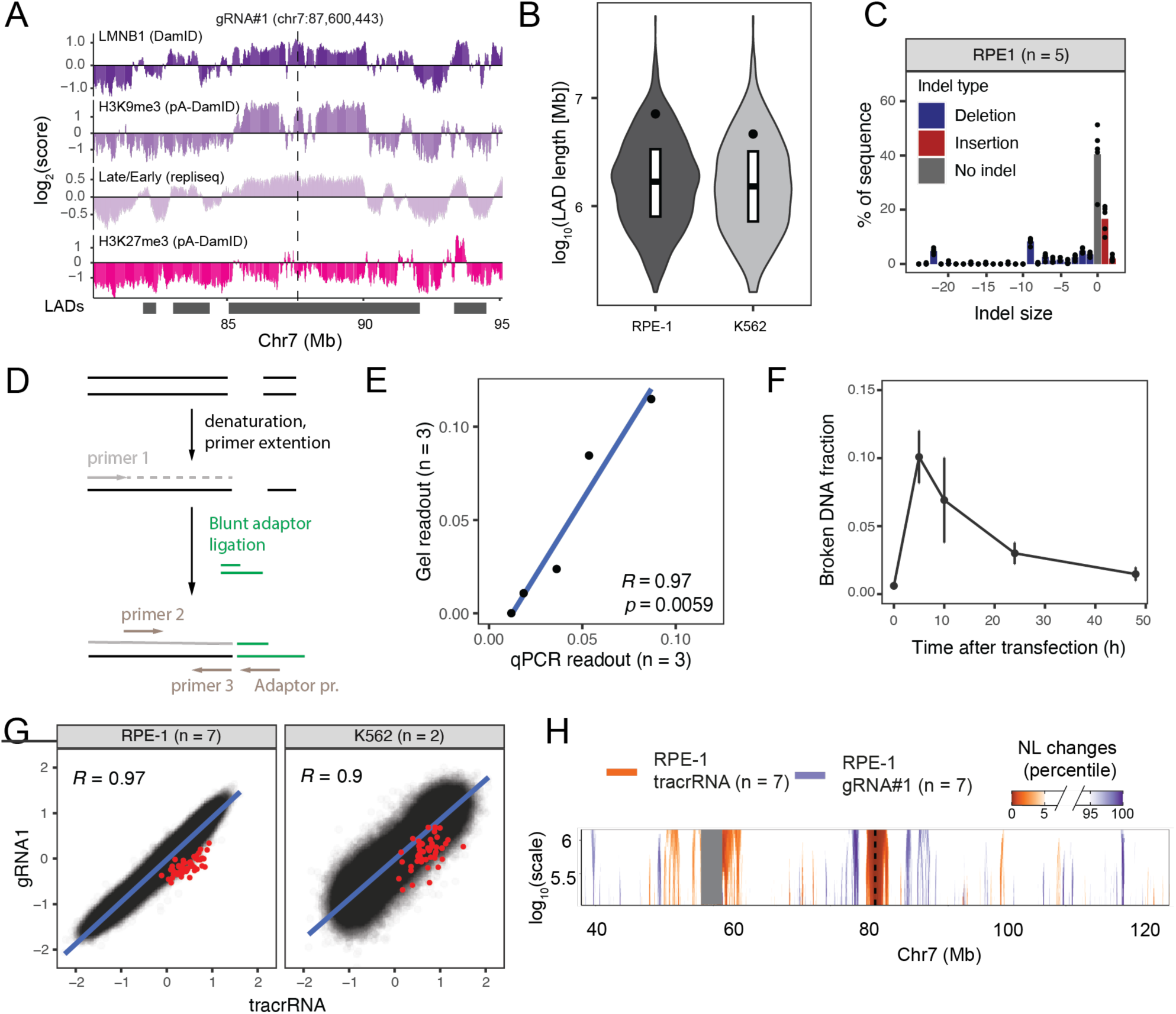
NL detachment in other cell types. (**A**) Heterochromatin features across a 15 Mb window (chr7:80.2-95.2 Mb) around the gRNA1 target site (vertical line) in RPE-1 cells. From top to bottom: log_2_(LMNB1/Dam) DamID track at 20 kb resolution from ^38^, log_2_(H3K9me3/Dam) pA-DamID track at 20 kb resolution from ^72^, log_2_(late/early) Repli-Seq track at 5 kb resolution from ^73^ and log_2_(H3K27me3/Dam) pA-DamID track at 20 kb resolution from ^72^. (**B**) Distribution of LAD lengths (Mb) in RPE-1 and K562 cells. The boxplot summarizes the 25^th^, 50^th^, and 75^th^ percentile of the distribution and the dot marks the length of LAD1. (**C**) Scheme of the main steps of the LM-PCR procedure and the primers used at each step. (**D**) Correlation between the broken DNA fraction measurements at different timepoints (n = 3) measured by qPCR (x-axis) and gel quantification (y-axis). The Pearson correlation coefficient and its corresponding p-value are shown in the bottom right corner. (**E**) Broken DNA fraction measurements at different timepoints after gRNA1 transfection. The dots represent the mean value over three biological and two technical replicates (**See methods**). The error bars represent the mean ± sd over the three biological replicates. (**F**) Indel percentage (%) after gRNA1 transfection in RPE-1 cells (n = 5) measured by TIDE. Each dot represents a biological replicate, and each bar is the mean percentage of each indel. The red bars show insertions, the blue ones show deletions, and the grey bar shows the percentage of sequence without indel. (**G**) Genome-wide correlation of NL interaction scores in tracrRNA only– and gRNA1-transfected RPE-1 (n = 7) and K562 cells at 20 kb resolution (n = 2). The plot shows the linear fit of the correlation as a blue line. The Pearson correlation coefficient is shown in the top-left corner. The red dots indicate the bins corresponding to a 1 Mb window around the break site (chr7:87.2-88.2 Mb). (**H**) Domainogram visualization, as described in Figure 1A, for a 100 Mb window around the gRNA1 target site in RPE-1 cells.

**Supplementary Figure 2:**
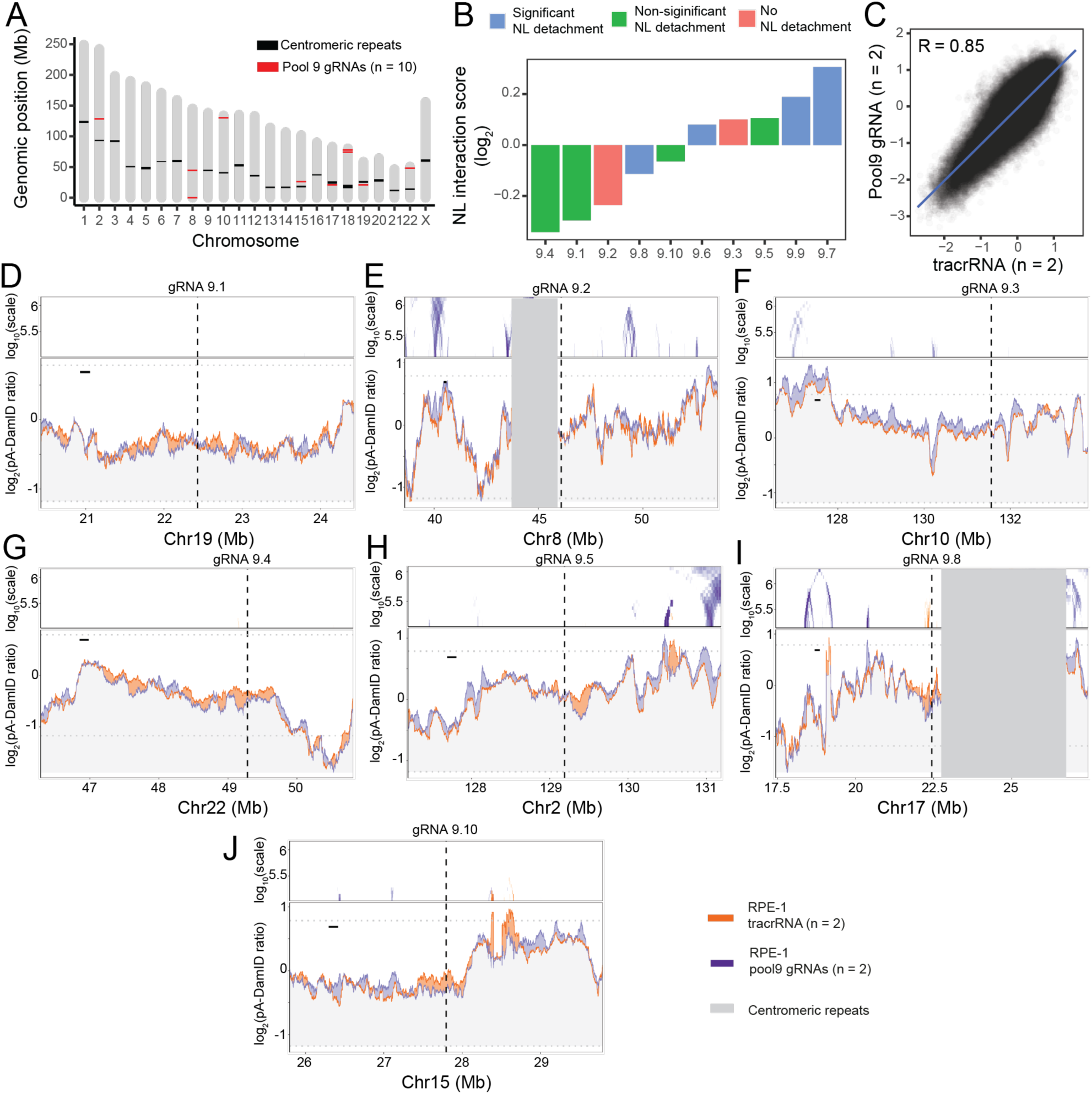
NL detachment occurs across multiple LADs in the genome. (**A**) Genomic coordinates of pool9 gRNA target sites (in red). Centromeric repeats of each chromosome are shown as black rectangles. (**B**) Mean NL interaction score (log_2_(LMNB2/Dam)) of a 1 Mb window around each gRNA target site in tracrRNA only-transfected RPE-1 cells. The gRNAs are ranked based on their NL interaction score (ascendent order), and their color indicates whether NL detachment is detected. (**C**) Genome-wide NL interaction score correlation of pool9– and tracrRNA only-transfected RPE-1 cells. The plot shows the data density as concentric circles, the fit of the linear regression in blue, and the Pearson correlation coefficient (R) in the top-left corner. NL interaction changes, visualized as described in Figure 1A, for pool9– and tracrRNA only-transfected RPE-1 cells. Each panel shows a different region of the genome: (**D**) 4 Mb around gRNA9.1 (chr19: 20.4-24.4 Mb), (**E**) 15 Mb around gRNA 9.2 (chr8: 38.5-53.5 Mb), (**F**) 7.6 Mb around gRNA9.3 (chr10: 126.55-133.79 Mb), (**G**) 4.5 Mb around gRNA9.4 (chr22: 46.28-50.81 Mb), (**H**) 4 Mb window around gRNA9.5 (chr2: 127.19-131.19 Mb), (**I**) 10 Mb window around gRNA 9.8 (chr17: 22.44-27.44 Mb), (**J**) 4 Mb window around gRNA 9.10 (chr15: 25.79-29.79 Mb). When centromeric repeats are within the visualized window, these are masked with a grey rectangle.

**Supplementary Figure 3:**
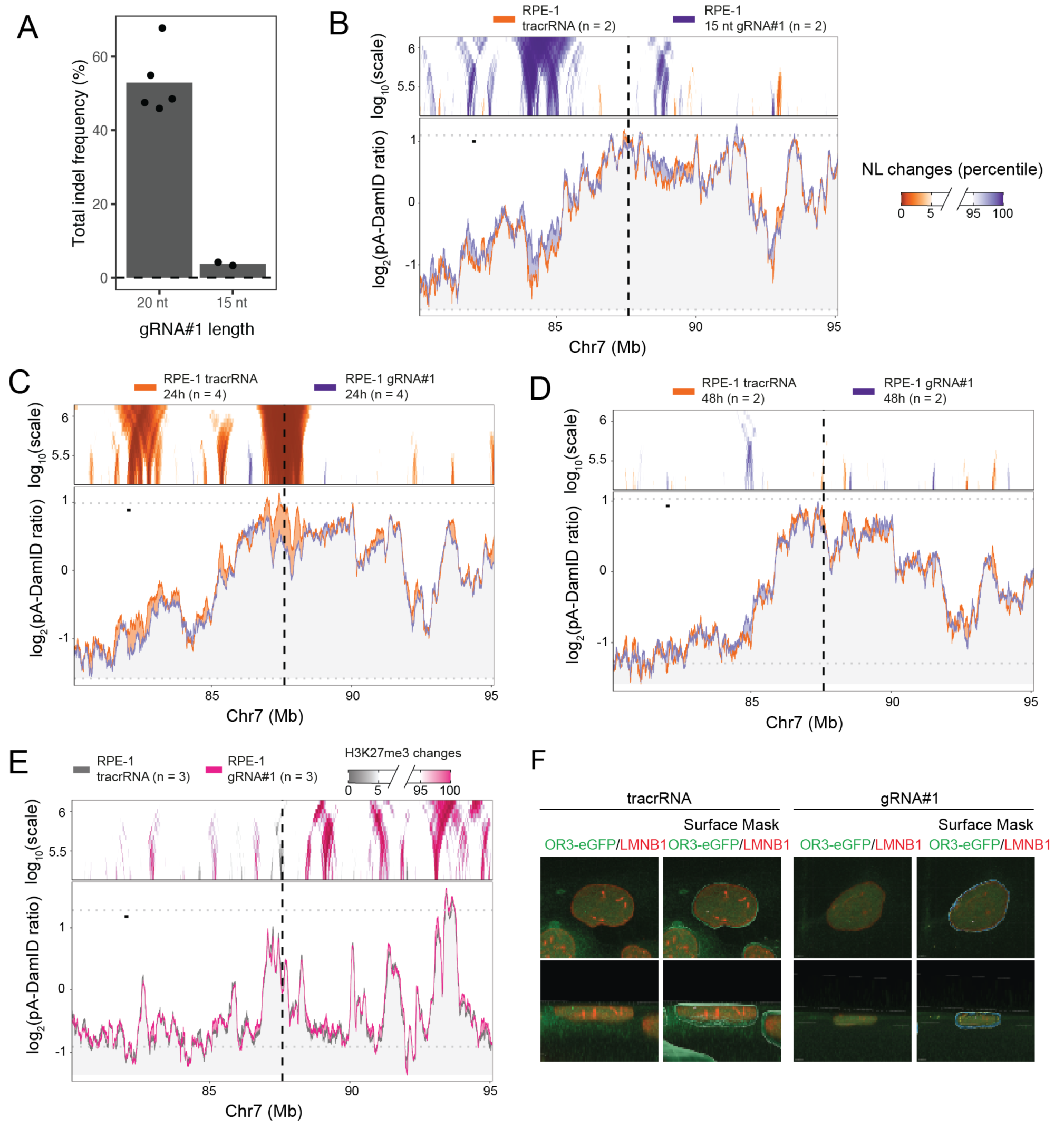
NL detachment is transient and dependent on formation of a DSB. (**A**) Total indel frequency (as percentages) measured by TIDE of RPE-1 cells transfected with the 20 nucleotide (nt) gRNA1 or a 15 nucleotide variant of gRNA1. Each dot represents a single replicate, and the grey bar is the mean over all replicates. (**B**) NL interaction changes after 15 nucleotide gRNA1 transfection (n = 2), plotted as described in Figure 1A. Parts of the genome that gain NL interaction are shown in purple and those that lose NL interaction are shown in orange. NL interaction changes (**C**) 24 hours and (**D**) 48 hours after gRNA1 transfection. The domainogram visualization is the same as in Figure 1A, and the rest of the plot shows pA-DamID interaction scores for tracrRNA only-(orange) and gRNA1-transfected (purple) RPE-1 cells. (**E**) H3K27me3 changes 10 hours after gRNA1 transfection, visualization as described in Figure 1A. Parts of the genome that gain H3K27me3 are shown in pink, and those that lose H3K27me3 are shown in grey. (**F**) Representative images of one 3D reconstructed ANCHOR3 RPE-1 cell after tracrRNA only (left) and gRNA1 (right) transfection. The images show the OR3-GFP signal in green and the LMNB1 (NL) signal in red. Each image in the top row shows an axial plane of one cell and in the bottom row a sagittal plane of one cell. For each condition the right show the NL surface mask in white and the OR3 focus in green.

**Supplementary Figure 4:**
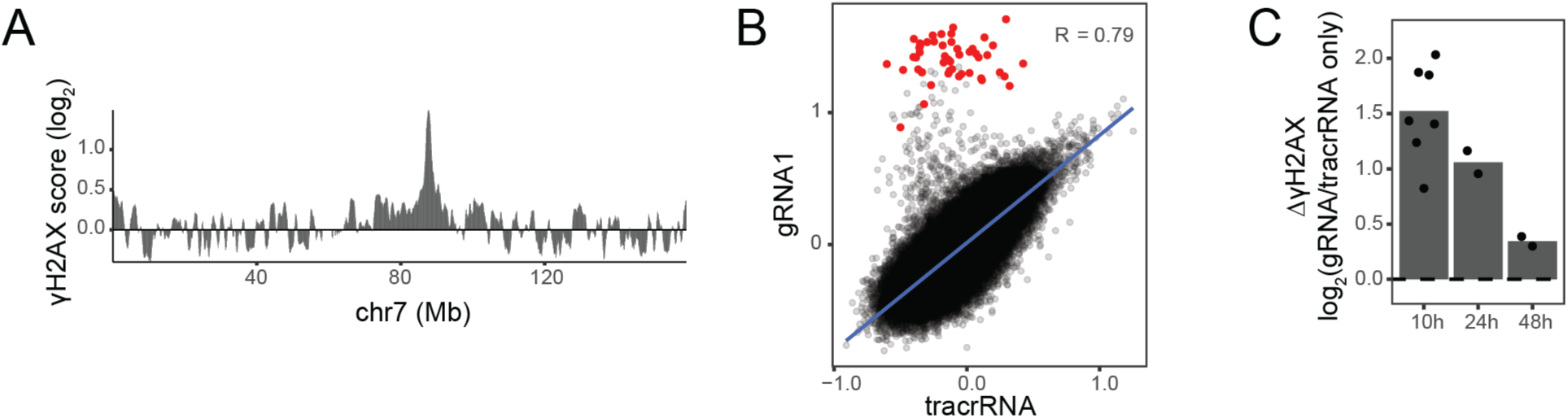
γH2AX accumulation correlates with NL detachment. (**A**) Whole chromosome log_2_(γH2AX/Dam) scores from gRNA1-transfected RPE-1 cells. (**B**) Same plot as in Figure S1G for gH2AX score in RPE-1 cells (n = 7). (C) ΔγH2AX scores, 10 (n = 7), 24 (n = 7), and 48 (n = 2) hours after gRNA1 transfection. The differential scores are calculated and visualized as those in Figure 1C.

**Supplementary Figure 5:**
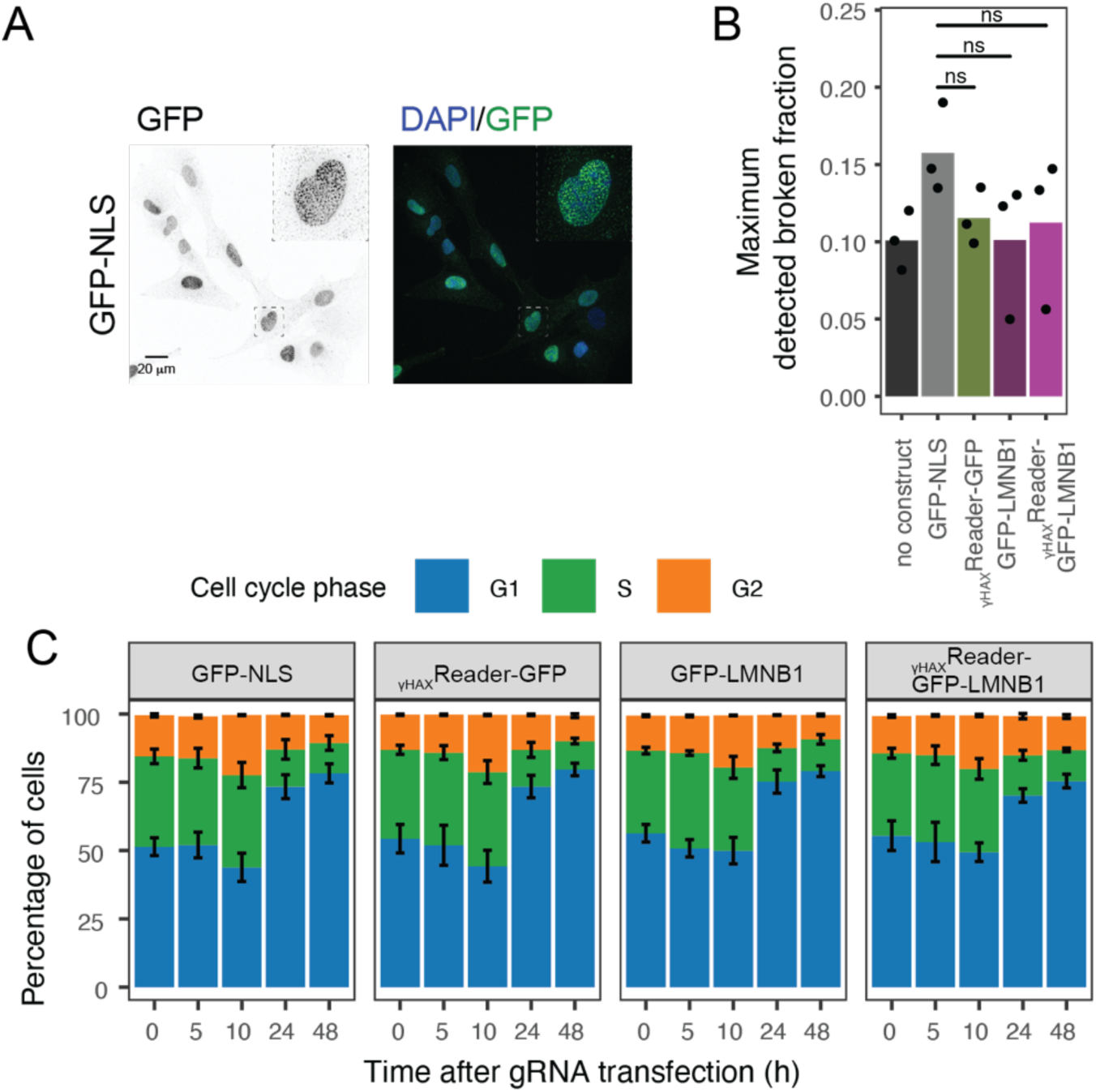
The expression of an NL tethering construct slows down repair. (**A**) Representative image of the GFP staining of RPE-1 cells overexpressing the GFP-NLS construct. From left to right, GFP immunofluorescence and composed image of both channels (DAPI in blue and GFP in green). (**B**) Maximum measured broken DNA fraction measured by LM-PCR in RPE-1 cells and cells expressing the NL-tethering construct or control constructs. Each dot represents a biological replicate (n = 3). The p-value is calculated using a two-sided Student’s t-test. (**C**) Cell cycle phase distribution of RPE-1 cells overexpressing the NL-tethering construct or control constructs at different timepoints after gRNA transfection. The bars show the mean percentage of cells in G1, S or G2 phases of the cell cycle. The error bars show the variability between the biological replicates as the mean ± sd values.

**Supplementary Figure 6:**
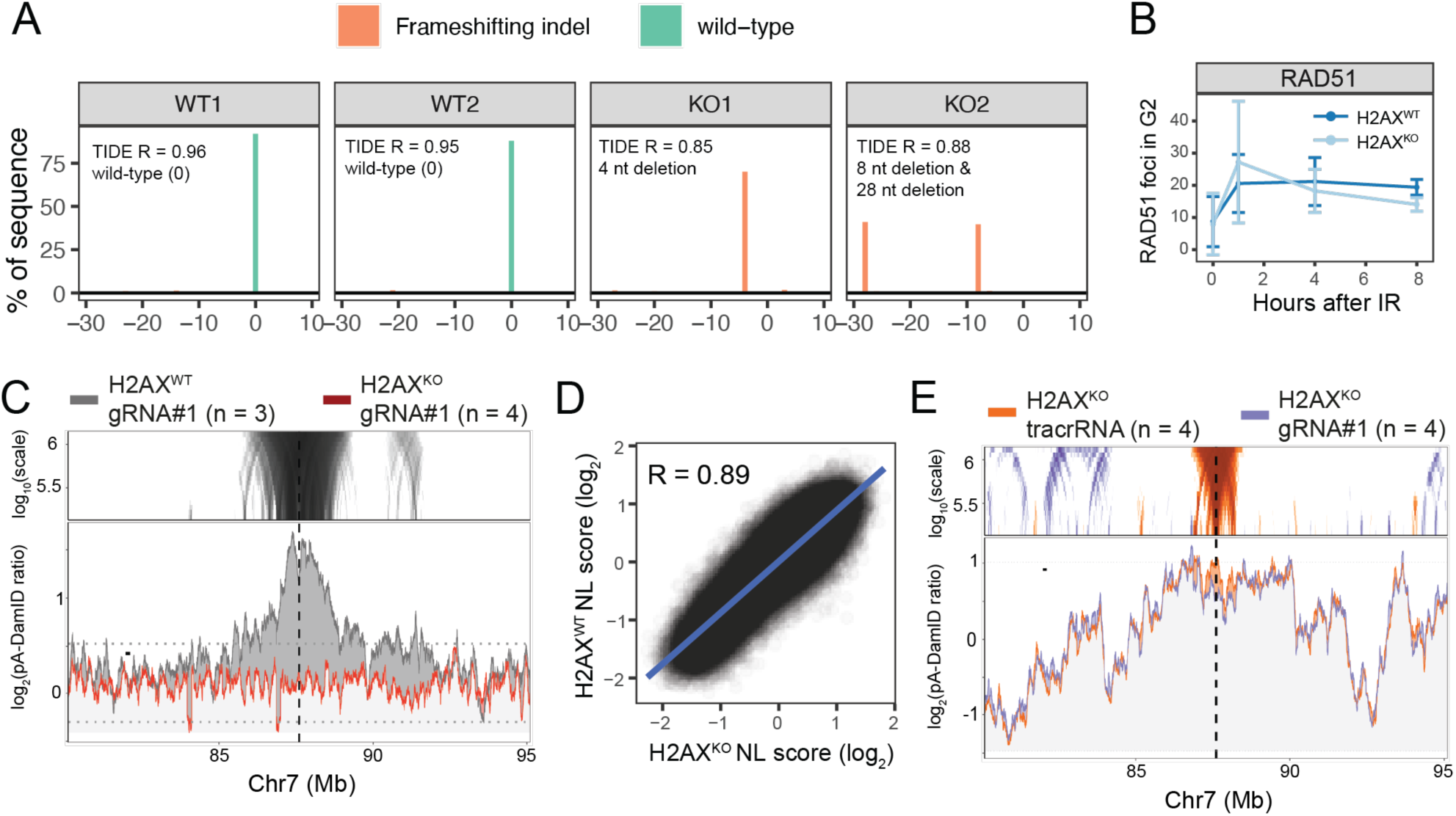
Characterization of H2AX^KO^ cells and NL detachment. (**A**) Genotyping of H2AX KO and WT clones by TIDE. The plots show the indel frequency around the targeted site in the first exon of the H2AFX gene. Each panel shows the sequence variation of a clonal cell line compared to the sequence in WT cells, and each bar represents a different indel with colors indicating whether the sequence variant is a wild-type sequence or a frameshifting mutation. In the top-left corner of each panel, the TIDE R coefficient and the main mutation for each clone are shown. (**B**) The mean number of RAD51 foci per G2 cell in H2AX^WT^ and H2AX^KO^ cells 1 hour (n = 6), 4 hours (n = 6), or 8 hours (n = 4) after 4 Gy irradiaton or no irradiation (0 hours, n = 6). The error bars show the mean ± sd values of all replicates. (**C**) γH2AX accumulation visualized as described in Figure 4A for H2AX^KO^ (red line) and H2AX^WT^ clones (grey) after gRNA1 transfection. The “specific” γH2AX domain was defined based on the (**D**) Genome-wide correlation of NL interaction (log_2_) scores in H2AX^WT^ (n = 3) and H2AX^KO^ (n = 4) clones. The plot shows the fit of the linear regression as a blue line, and Pearson’s R. (**E**) NL interaction changes in H2AX^KO^ cell transfected with gRNA1 or tracrRNA only visualized as described in Figure 1C. The parts of the genome highlighted in orange show the parts of the genome that lose NL interaction after gRNA1 transfection in H2AX^KO^ cells.

**Supplementary Figure 7:**
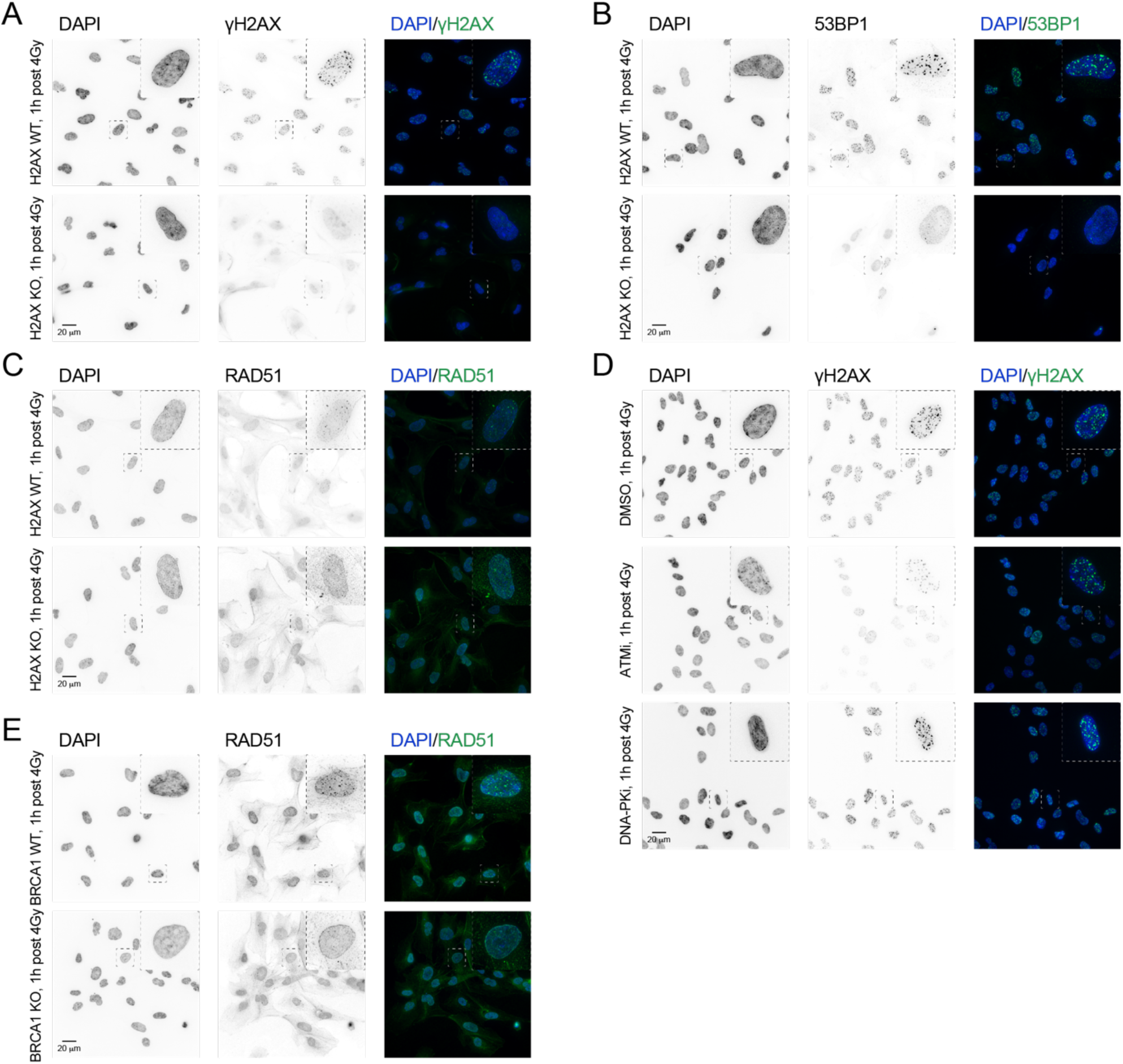
Representative images of γH2AX, 53BP1 and Rad51 foci. (**A**) Representative image of the γH2AX staining of a H2AX^WT^ (top) and H2AX^KO^ (bottom) cell 1 hour after 4Gy irradiation. From left to right, DAPI staining showing the nucleus of the cell, γH2AX immunofluorescence staining and composed image of both channels (DAPI in blue and γH2AX in green. (**B**) Same as **Figure S6A** for 53BP1 staining. (**C**) Representative image of the γH2AX staining of an RPE-1 cell treated with vehicle-only (DMSO, top), ATM (middle) or DNAPK inhibitor (bottom) 1 hour after 4Gy irradiation. From left to right, DAPI staining showing the nucleus of the cell, γH2AX immunofluorescence staining and composed image of both channels (DAPI in blue and γH2AX in green. (**D**) Representative image of the RAD51 staining of an p53^KO^ (BRCA1 WT) and p53/BRCA1^dKO^ (BRCA1 KO) cell 1 hour after 4Gy irradiation. From left to right, DAPI staining showing the nucleus of the cell, RAD51 immunofluorescence staining and composed image of both channels (DAPI in blue and RAD51 in green).

**Supplementary Figure 8:**
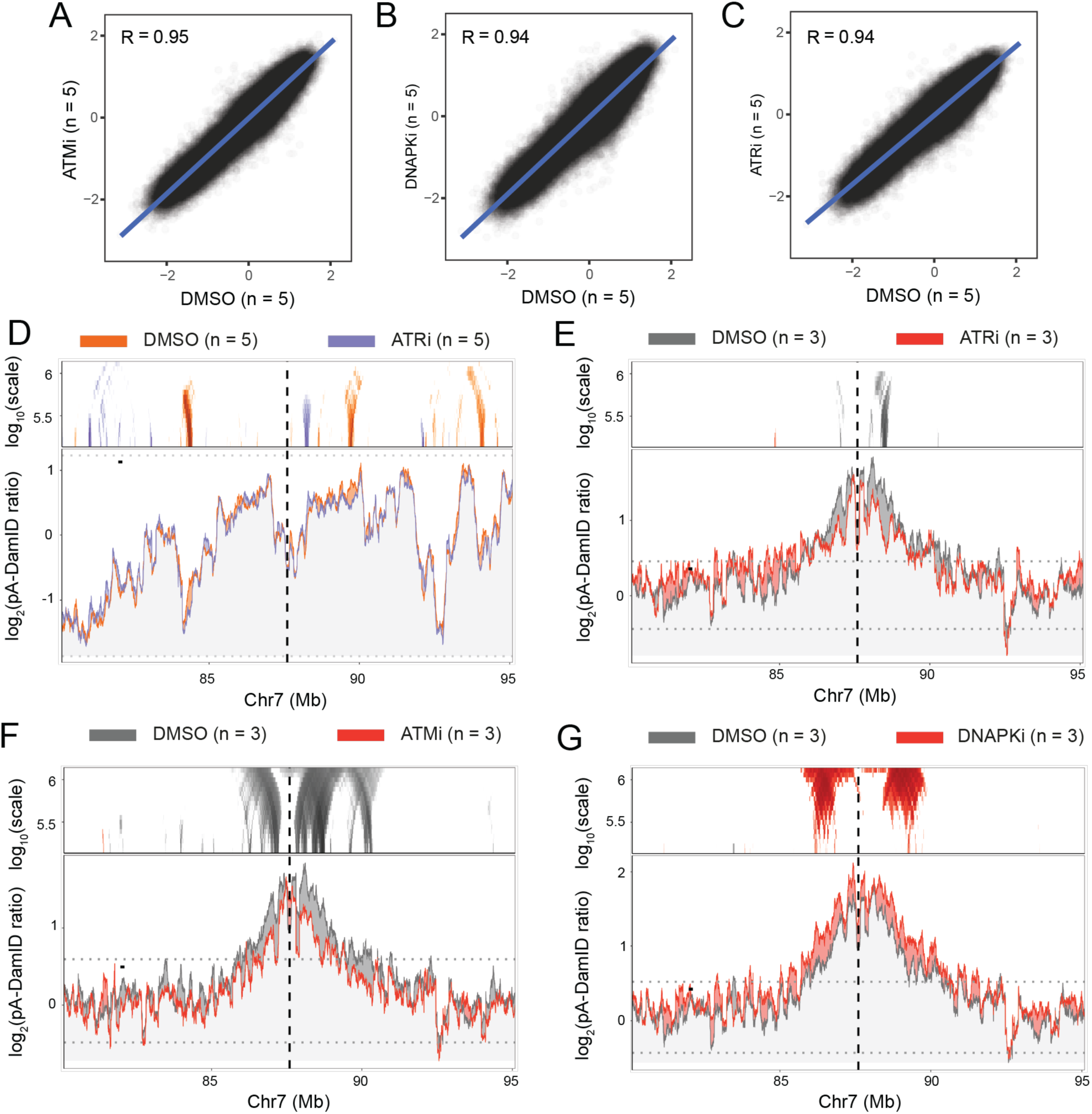
NL interaction changes in RPE-1 cells treated with inhibitors for ATM, DNAPK_cs_, or ATR. Genome-wide correlation of NL interaction (log_2_) scores in inhibitor or vehicle-only treated RPE-1 cells (n = 5) for (**A**) ATM inhibitor, (**B**) DNAPK_cs_ inhibitor, and (**C**) ATR inhibitor. The plot shows the fit of the linear regression as a blue line, and Pearson’s R. (**D**) NL interaction (n = 5) changes in ATR inhibitor-treated and gRNA1-transfected RPE-1 cells visualized as previously described in Figure 1A. γH2AX changes in (**E**) ATR inhibitor-(**F**) ATM inhibitor-(n = 3), and (**G**) DNAPK_cs_ inhibitor-treated (n = 3) and gRNA1-transfected RPE-1 cells. These data are visualized as described in Figure 3A.

**Supplementary Figure 9:**
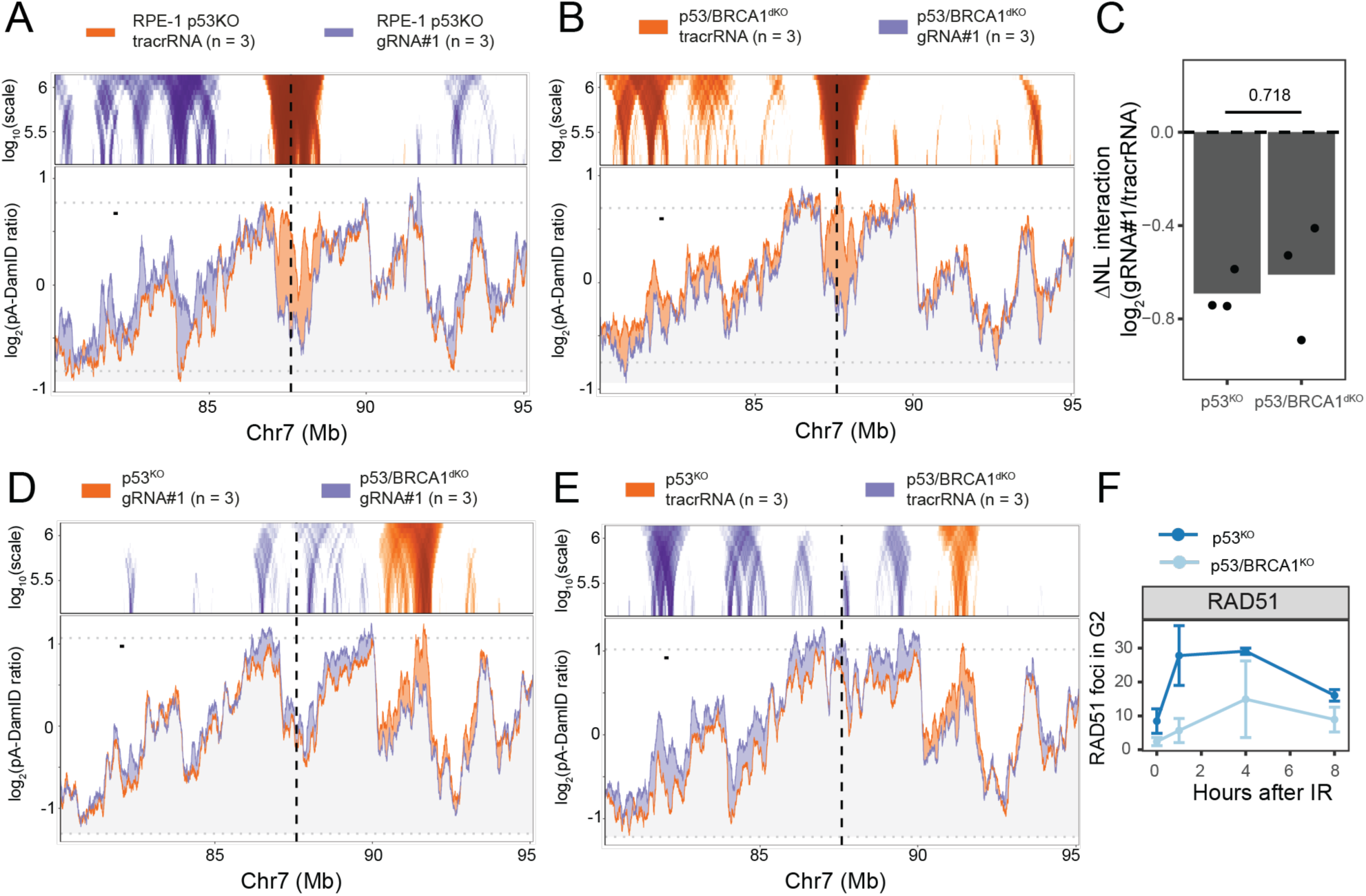
NL detachment does not depend on BRCA1. (**A**) NL interaction changes in RPE-1 p53^KO^ cells 10 hours after tracrRNA only (n = 3) or gRNA1 (n = 3) transfection. The changes are visualized as described in Figure 1A. In brief, parts of the genome that gain NL interaction are shown in purple, and parts of the genome that lose NL interaction are shown in orange. (**B**) Same as in **Figure S8A** but for RPE-1 p53/BRCA1^dKO^ cells. (**C**) Differential NL interactions after gRNA1 transfection in p53^KO^ and p53/BRCA1^dKO^ cells. The ΔNL interaction is calculated as in Figure 1C, each dot represents a single replicate, and the p-value is calculated using a two-sided paired Student’s t-test. Same visualization as in Figure 5C for NL interaction changes between (**D**) gRNA1– and (**E**) tracrRNA only-transfected p53^KO^ (n = 3, control) and p53/BRCA1^dKO^ RPE-1 cells (n = 3, experimental). (**F**) Number of RAD51 foci per G2 cell at different time points after 4Gy irradiation (1 (n = 3), 4 (n = 3) and 8 (n = 2) hours) or without irradiation (0 hours, n = 3) in RPE-1 p53^KO^ or p53/BRCA1^dKO^ cells. Each dot shows the mean number of foci of all replicates, and the error bars show the mean ± sd values.

**Supplementary Figure 10:**
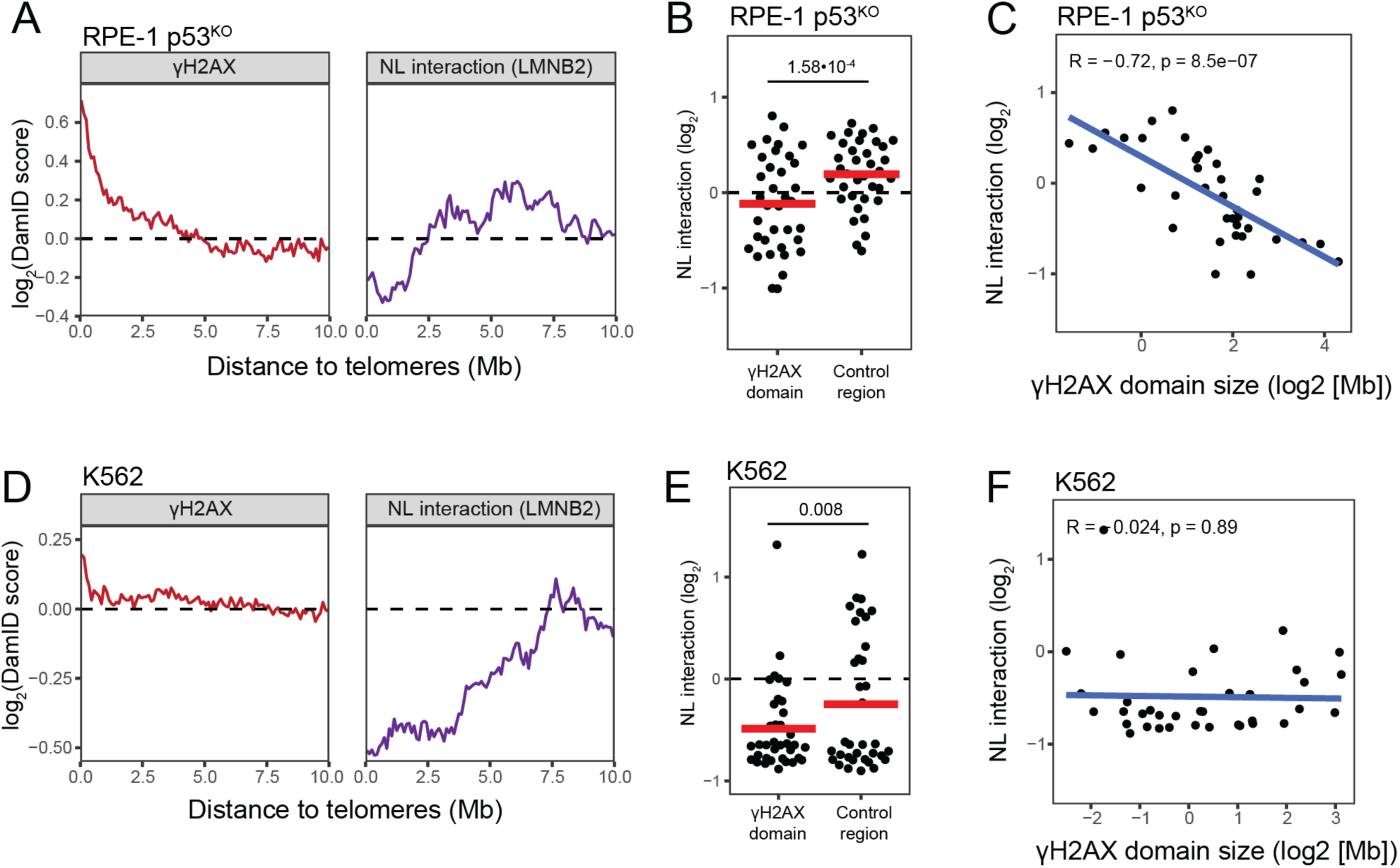
gH2AX accumulation at chromosome ends inversely correlates with NL association in additional cell lines. Comparison between gH2AX and NL interaction scores of chromosome ends in (**A-C, top**) RPE-1 p53^KO^ and (D-F, bottom) K562 cells. Each row consists of three plots: (**A&D**) Mean gH2AX (left) and NL interaction (right) scores as a function of distance to the telomeres. The same plot as in Figure 6D. (**B&E**) Mean NL interaction scores (log2) of subtelomeric gH2AX domains and adjacent control regions. Each dot represents a single chromosome end as described in Figure 6E. (**C&F**) Correlation plot between NL interaction scores (log2) and gH2AX domain size (Mb). The same analysis and graph as in Figure 6F.

## SUPPLEMENTARY TABLE

**Supplementary Table 1:**
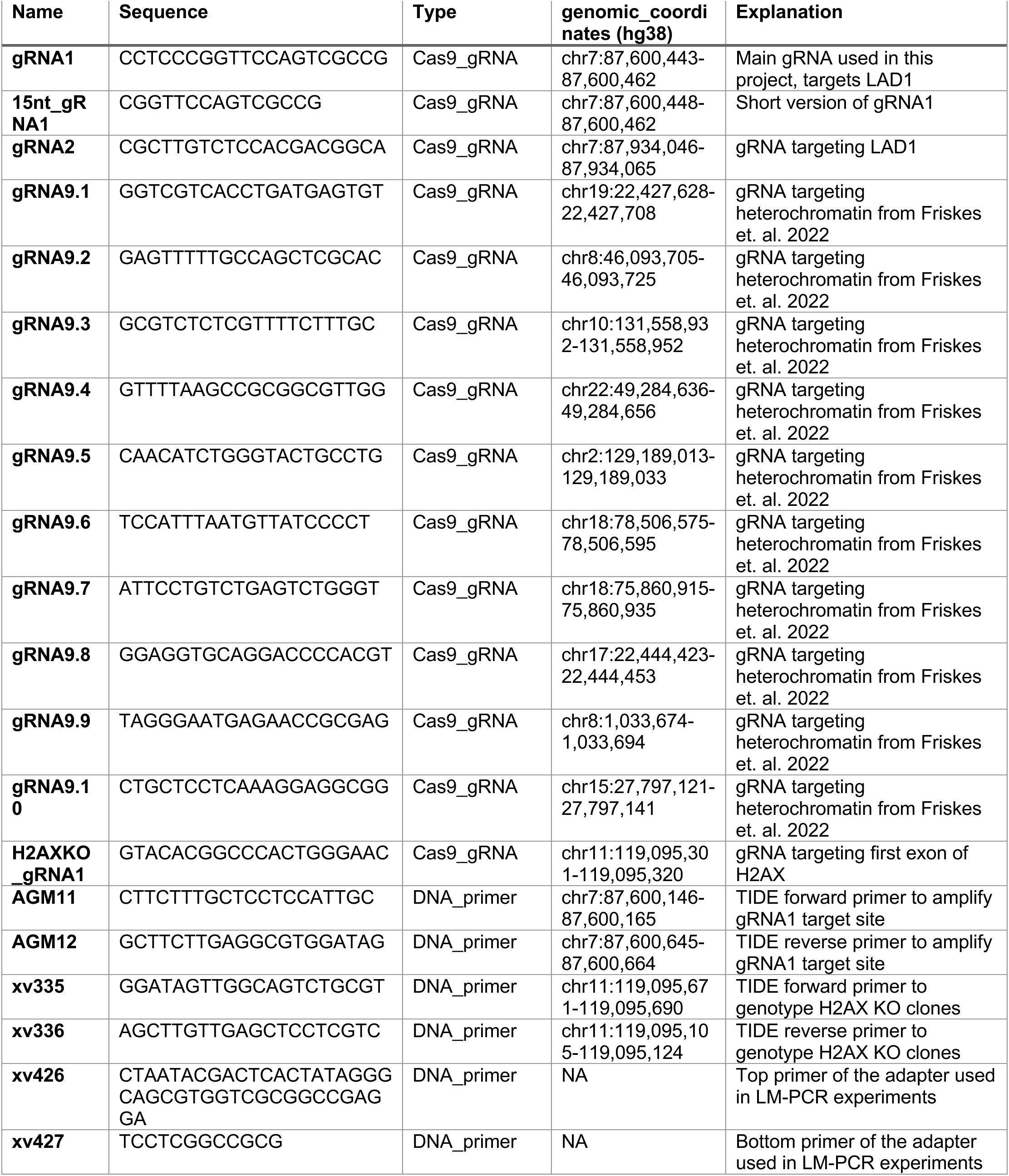
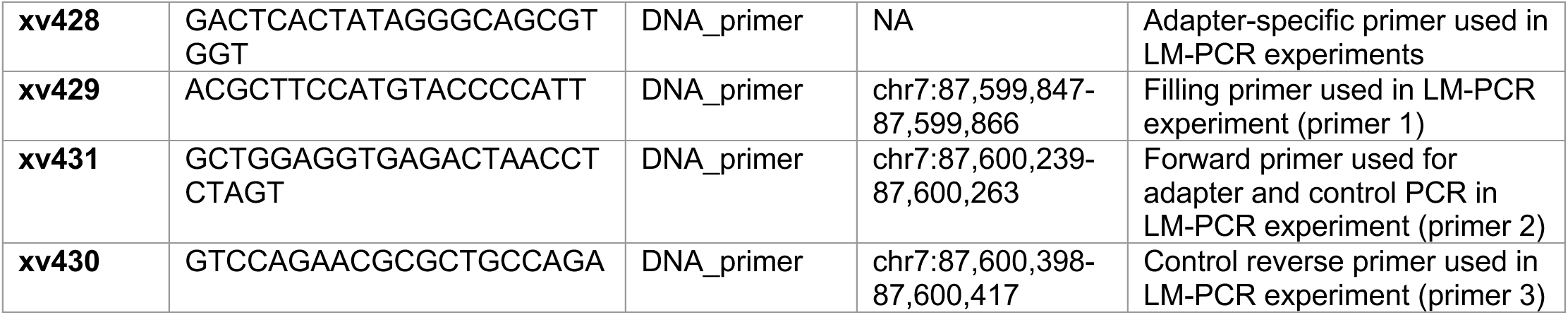
Oligonucleotide sequences used in this study.

